# Human and murine *Cryptococcus neoformans* infection selects for common genomic changes in an environmental isolate

**DOI:** 10.1101/2022.04.12.487930

**Authors:** Poppy Sephton-Clark, Scott A. McConnell, Nina Grossman, Rosanna Baker, Quigly Dragotakes, Yunfan Fan, Man Shun Fu, Gracen Gerbig, Seth Greengo, J. Marie Hardwick, Madhura Kulkarni, Stuart M. Levitz, Joshua D. Nosanchuk, Shmuel Shoham, Daniel Smith, Piotr Stempinski, Maggie Wear, Christina A. Cuomo, Arturo Casadevall

## Abstract

A pet cockatoo was the suspected source of *Cryptococcus neoformans* recovered from the cerebral spinal fluid (CSF) of an immunocompromised patient with cryptococcosis based on the molecular analyses available in 2000. Here we report whole genome sequence analysis of the clinical and cockatoo strains. Both are closely related MATα strains belonging to the VNII lineage, confirming that the human infection likely originated from pet bird exposure. The two strains differ by 61 single nucleotide polymorphisms, including 8 nonsynonymous changes involving 7 genes. To ascertain whether changes in these genes are selected during mammalian infection, we passaged the cockatoo strain in mice. Remarkably, isolates obtained from mouse tissue possess a frame-shift mutation in one of the seven genes altered in the human sample, a gene predicted to encode a SWI-SNF chromatin-remodeling complex protein. Both cockatoo and patient strains as well as mouse passaged isolates obtained from brain tissue had a premature stop codon in a homolog of ZFC3, a predicted single-zinc finger containing protein, which is associated with larger capsules when deleted and appears to have reverted to a full-length protein in the mouse passaged isolates obtained from lung tissue. The patient strain and mouse passaged isolates show variability in the expression of virulence factors, with differences in capsule size, melanization, and rates on non-lytic expulsion from macrophages observed. Our results establish that environmental strains undergo genomic and phenotypic changes during mammalian passage, suggesting that animal virulence can be a mechanism for genetic change and that the genomes of clinical isolates may provide a readout of mutations acquired during infection.

## Introduction

*Cryptococcus neoformans* is a human pathogenic fungus that is a major cause of life-threatening meningoencephalitis (1). Cryptococcosis is more common in patients with impaired immune systems although occasional disease occurs in individuals with no apparent immune deficits. *C. neoformans* infection is first thought to be acquired in childhood (2) and is either cleared or can become latent to reactivate if impaired immunity occurs later in life (3). However, disease can also follow exposure to contaminated environmental sources in adults, but the ubiquity of this fungus complicates the identification of point sources. Restriction enzyme polymorphism analysis of patient and environmental samples in New York City revealed that some clinical isolates shared the same restriction patterns and were thus indistinguishable from local infection sources (4), but such analysis lacked the precision to reveal point sources, particularly given that the disease often develops slowly and can be the result of latent, distantly acquired, infection (5). A recent investigation of a cryptococcosis outbreak in a Scottish hospital revealed how difficult it is to make associations between clinical, and geographically and temporally matched environmental samples (6).

In 2000, we reported on the case of an immunosuppressed patient with cryptococcosis who had a pet cockatoo (7). *Cryptococcus* was also recovered from the cockatoo guano, which was not unexpected as bird guano is a common environmental reservoir for *C. neoformans* (8, 9). Both the patient and bird guano strains were indistinguishable using molecular tools available at the time, resulting in the first example of cryptococcal infection traced from a point source (7). In subsequent years additional cases of *C. neoformans* infections linked to pet birds were reported (10, 11). Since the original report, genomic sequencing has become routine and there is now a wealth of information available on *C. neoformans* genomes (12). In this study, we compared the genome sequences of the patient and cockatoo strains and passaged the cockatoo strain in mice to identify genetic and phenotypic changes resulting from mammalian infection. The results validate the earlier conclusion that the clinical and cockatoo strain sequences were closely related, and that the cockatoo was the likely source of infection. We also find evidence of similar genome evolution in mouse-passaged and patient strains. These results implicate novel genomic loci in virulence, identify functionally similar genomic changes arising from mammalian infection, and characterize phenotypic differences between mouse passaged isolates that appear to be organ specific.

## Materials and Methods

### Resource Availability

Requests for strains used in this study should be directed to Arturo Casadevall.

### Experimental model and subject details

#### *C. neoformans* strains

The original patient and cockatoo strains were described in Nosanchuk et al., (7) and have been maintained frozen at -80 °C since last studied. This study involved the analysis of original clinical and cockatoo strains as well as mouse passaged cockatoo isolates.

#### Mouse passage studies

An inoculum of 2 × 10^5^ freshly grown *C. neoformans* (CU) cells from the cryopreserved samples obtained from cockatoo guano was administered by retroorbital intravenous injection to a female A/J mouse. Infections were administered to the mouse under xylazine and ketamine anesthesia, 10 mg/kg and 100 mg/kg, respectively. The mouse was observed daily for signs of cryptococcosis symptoms and eventually euthanized 43 days post-infection. Brain and lungs were aseptically removed and homogenized in a total volume of 2 mL sterile PBS and dilutions of 10^−4^, 10^−3^, 10^−2^, 10^−1^, and 10° were plated onto YPD agar plates supplemented with 1% Pen/Strep (Gibco 15140122) to determine CFUs in each organ. The brain and lung homogenates contained 8.16 × 10^2^ and 7.88 × 10^1^ CFU/mg, respectively.

#### Animal studies

An A/J female mouse 5-8 wk of age was obtained from The Jackson Laboratory (Bar Harbor, ME). Animal experiments were conducted in accordance with the policies and with the approval of the Johns Hopkins University Institutional Animal Care and Use Committee (protocol MO21H124). Mice were sacrificed using CO2 asphyxiation.

#### Macrophage non-lytic event quantification

Bone marrow derived murine macrophages (BMDMs) were harvested from the hind leg bones of 6-week-old C57BL/6 female mice from The Jackson Laboratory and were differentiated by seeding in 10 cm tissue culture treated dishes in DMEM with 10% FBS, 1% nonessential amino acids, 1% penicillin-streptomycin, 2 mM Glutamax, 1% HEPES buffer, 20% L-929 cell conditioned supernatant, and 0.1% beta-mercaptoethanol for 6 days at 37 °C and 9.5% CO_2_. BMDMs were used for experiments within 5 days of differentiation. BMDMs were activated with LPS (0.5 ug/mL) and IFN-γ (10 ng/mL) for 16 h prior to experiments. The media was then refreshed and the BMDMs were infected with opsonized *C. neoformans* at MOI 1, then imaged every 2 min for 24 h on a Zeiss Axiovert 200M inverted scope at 37 °C with 9.5% CO_2_. Non-lytic events were quantified by determining the outcome of each infected macrophage throughout the 24-h period and 95% confidence intervals were estimated with a test of proportions. Statistical significance was determined with a test of equal proportions and Bonferroni correction.

#### Amoeba predation analysis

A modified version of the fungal killing assay described previously (26) was used to determine the ability of different fungal strains to resist killing by *Acanthamoeba castellanii*. Briefly, *A. castellanii* were washed twice with Dulbecco’s phosphate buffered solution (DPBS) supplemented with calcium and magnesium, counted in a hemocytometer and diluted with DPBS to 1 × 10^4^ cells/ml. Amoeba were seeded in 96 well plates and incubated at 25°C for one hour to allow for adherence to the bottom of the plate. *C. neoformans* cultures were grown overnight in liquid yeast peptone dextrose (YPD) at 30°C, washed twice with DPBS, and similarly counted and diluted in DPBS. Wells containing amoebae and control wells containing only DPBS were inoculated with *C. neoformans* at 1 × 10^4^ cells/well and incubated at 25°C for 0, 24, or 48 h. Amoebae were lysed by passage through a 27-gauge needle seven times and then lysates serially diluted. Three 10 µL samples of each the serially diluted lysates were plated on YPD agar and incubated at 30°C for 48 h. CFUs were counted with a light microscope. Total CFUs were calculated, and significance was determined with two-way ANOVA.

#### Virulence in *Galleria mellonella*

*G. mellonella* were infected with CU, PU, CPB, and CBL isolates as previously described (3). Briefly, final instar larvae ranging between approximately 175 and 225 mg were injected with 10 µl of *C. neoformans* culture grown to stationary phase in YPD media, washed twice in PBS, and diluted to a concentration of 10^7^ cells/ml. Survival of larvae and pupae was monitored daily for 14 days, with death being defined by lack of movement in response to the stimulus of a pipette tip. Statistical significance was determined with the log-rank Mantel-Cox test, corrected for multiple comparisons using the Bonferroni method, via GraphPad Prism.

## Method Details

### India ink staining and capsule analysis

*C. neoformans* cells were mixed with India ink and imaged on an Olympus AX70 microscope using QImaging Retiga 1300 digital camera and the QCapture Suite V2.46 software (QImaging). Capsule measurements were calculated using the exclusion zone produced with India ink and the Quantitative Capture Analysis program as previously described (13). A minimum of 100 yeast cells were measured for each strain and condition. Statistical significance was determined with paired two-tailed T-tests with unequal variance, via Microsoft Excel.

### Growth curves

*C. neoformans* strains were recovered from cryopreserved stocks by growth in YPD for 48 h at 30°C and then sub-cultured in triplicate into Sabouraud dextrose broth (BD Difco) at a density of 1 × 10^5^ cells/mL in wells of a 96-well plate. The plate was incubated at 30 or 37°C with orbital shaking in a SpectraMax iD5 plate reader (Molecular Devices) and absorbance at 600 nm was read at 15 min intervals. Absorbance was plotted against time using GraphPad Prism software and the linear region of each curve was analyzed by simple linear regression to derive the slope and lag time (x-intercept).

### Exopolysaccharide preparation

*C. neoformans* CU, PU, CPB, and CPL isolates were grown in YPD for 48 h and sub-cultured to modified minimal media (MMM) for NMR analysis which also induces capsule growth. Cells were grown in MMM for 3 days and the exo-polysaccharide (EPS) was harvested by centrifuging cells (10 minutes at 3494 x g) and sterile filtration of culture supernatant (0.45 nm filtration). Any remaining media components were removed, and EPS was concentrated by passage through a 3 kDa molecular weight cut-off (MWCO) centricon filtration unit. The > 3 kDa fraction of EPS was then characterized by NMR.

### NMR analysis

1D ^1^H NMR data were collected on a Bruker Avance II (600 MHz), equipped with a triple resonance, TCI cryogenic probe, and z-axis pulsed field gradients. Spectra were collected at 60°C, with 128 scans and a free induction decay size of 84336 points. Standard Bruker pulse sequences were used to collect the 1D data (p3919gp and zggpw5). Data were processed in TopSpin (Bruker version 4.1.3) by truncating the FID to 8192 points, apodizing with a squared cosine bell window function and zero filling to 65536 points. Relative peak integration and ^1^H frequency referencing was performed on TopSpin using d_6_-DSS internal standard (10µl/510 µl sample).

### DNA preparation and genomic sequencing

Oxford Nanopore Technologies (ONT) sequencing libraries were prepared with genomic DNA from the CU and PU strains using the Ligation Sequencing Kit (SQK-LSK109) with the Native Barcoding Kit (EXP-NBD103) according to manufacturer instructions (Oxford Nanopore Technologies, Oxford, UK). Illumina sequencing libraries were prepared using the Nextera Flex DNA library prep kit (Illumina, San Diego, California) and sequencing was performed with a MiSeq using v2 2×150 chemistry. For genomic sequencing of CPL and CPB isolates, three individual colonies were selected from brain and lung CFU plates and used to seed 100 mL YPD cultures, which were allowed to grow for 48 h at 30°C with rotation. Genomic DNA was isolated from each culture following the protocol described in Velegraki et al. (14). Briefly, after a 48 h growth period, *C. neoformans* cells were collected by centrifugation, frozen at -80°C overnight, then subsequently lyophilized overnight. Glass beads (0.5 mm diameter, BioSpec Products, Cat No. 11079105z) were added to the dry cell pellets and vortexed into a powder. DNA was then extracted with a CTAB buffer (100 mM Tris, pH 7.5, 700 mM NaCl, 10 mM EDTA, 1% CTAB, 1% beta-mercaptoethanol) at 65°C for 30 min. The tubes were cooled and then extracted with chloroform, then isopropanol, then 70% ethanol. The DNA pellet was then resuspended in 1 mL sterile water and treated with 20 µg RNase for 30 min at 37°C. Finally, the genomic DNA was further purified using the DNeasy PowerClean CleanUp Kit (Qiagen 12877-50) for genomic sequencing. 100 uL aliquots of 19-30 ng/µL DNA were used in the sequencing reactions. DNA was sheared to 250bp using a Covaris LE instrument and adapted for Illumina sequencing as described by Fisher et al. (15). Libraries were sequenced on a HiSeq X10, generating 150bp paired reads.

### Assembly and genomic analysis

Whole genome assemblies were generated for CU and PU strains with ONT long reads via Canu v2.1.1 (genome size 20Mb) (16), followed by short read polishing via medaka v0.8.1 (1X) (https://github.com/nanoporetech/medaka) and pilon v1.23 (3X)(17). Single nucleotide polymorphisms, insertions, and deletions between CU and PU assemblies were then identified using nucmer v3.1 (18), with variants in centromeric and telomeric regions removed prior to downstream analysis. To identify variants in the mouse passaged isolates, Illumina reads for CPL and CPB samples were aligned to the CU assembly with BWA-MEM v0.7.17 (19), and variants were called with our publicly available GATK v4 pipeline (https://github.com/broadinstitute/fungal-wdl/tree/master/gatk4). Post calling, variants were filtered on the following parameters: QD < 2.0, QUAL < 30.0, SOR > 3.0, FS > 60.0 (indels > 200), MQ < 40.0, GQ < 50, alternate allele percentage = 0.8, DP < 10. All variants were annotated with SNPeff, v4.3t (20). To identify strain lineage, reads for the CU and PU samples were aligned to the *Cryptococcus neoformans var. grubii* H99 reference genome (GCA_000149245.3) with BWA-MEM v0.7.17 (19), and variants were called and filtered as described above. A maximum likelihood phylogeny was estimated using segregating SNP sites present in one or more isolates, allowing ambiguity in a maximum of 10% of samples, with RAxML v8.2.12 (21) rapid bootstrapping (GTRCAT substitution model), and visualized with ggtree (R 3.6.0) (22). Aneuploidies were visualized using funpipe (coverage analysis) v0.1.0 (https://github.com/broadinstitute/funpipe), transposon mobilization was assessed through whole genome alignment of the CU and PU assemblies with nucmer v3.1 to identify alignment gaps, and copy number variation was assessed using CNVnator v0.3 (23).

### Urease activity assay

*C. neoformans* strains and isolates were first grown in YPD for 48 h at 30°C. Urea broth comprised of 10 mM KH_2_PO_4_, 0.1% Bacto Peptone (Difco), 0.1% D-glucose, 0.5% NaCl, 2% urea, and 0.03 mM phenol red, as described by Roberts et al., (24), was inoculated with PBS-washed cells at a density of 1 × 10^6^ cells/mL for each strain in triplicate. After incubation for 16 h at 30°C, increased pH of culture media that is indicative of ammonium production due to urease activity was detected by measuring absorbance at 560 nm over a 6 h time course. Absorbance readings that had been corrected by subtraction of media-only background absorbance were plotted against time. Urease activity rates were derived by a simple linear regression model and data were analyzed for statistical significance using an ordinary one-way analysis of variance (ANOVA) with GraphPad Prism 9 software.

### *C. neoformans* melanization assay

*C. neoformans* strains and isolates were first grown in YPD liquid media for 48 h at 30°C until the cultures were in stationary phase. Cells were washed twice in PBS. 100 µl of washed culture was added to 5 mL Minimal Media with 1 mM L-3,4-dihydroxyphenylalanine (L-DOPA) and grown for 5 d at 30°C. Cultures were removed and placed into a 6-well plate for imaging. Alternatively, 1 × 10^6^ PBS-washed cells were spotted onto L-DOPA agar in triplicate. Plates were incubated at 30°C or 37°C and then photographed after 2, 3, and 6 days using a 12-megapixel camera. Color images were converted to grayscale using Adobe Photoshop, pigmentation intensities were quantified using Image Studio Lite software, and graphed using GraphPad Prism software. Statistical significance was determined with the ordinary one-way ANOVA test, via GraphPad Prism.

### Phospholipase activity

Extracellular phospholipase in *C. neoformans* strains and isolates was tested by the modified method reported by the Chen et al. (25). Egg yolk agar medium was created based on Difco Sabouraud Dextrose Agar media with 8% egg yolk, 1M sodium chloride and 0.05M calcium chloride. After overnight growth in YPD media 3 ul of each strain, with a total of 10,000 cells, were spotted onto egg yolk agar medium. Each strain and isolate was tested on five separate plates and incubated at 30°C. After 72 h and 96 h colonies were photographed and measured with ImageJ software. Activity of phospholipase was analyzed by the ratio of colony diameter to total precipitation diameter, where a ratio equal to 1.0 indicates a lack of phospholipase activity. Statistical significance was determined by an unpaired t-test test, via GraphPad Prism.

### Heat-ramp and thermal stability analysis

Strains and isolates were maintained at -80°C in glycerol, streaked onto Sabouraud dextrose (SAB) (BD Difco) agar and incubated at 30°C for 48 h prior to heat-ramp cell death assays. SAB broth was inoculated from growth on plates to equal densities (OD 0.1) and incubated at 30°C for 18 h in stationary 96-well plates. Isolates were resuspended, diluted 1:5 in fresh SAB broth, and 100 µl was treated with a linear 30°C to 56°C heat-ramp stress over 10 min in a water bath with agitation (Lauda). Untreated and heat-ramp treated strains were immediately spotted (5 µl in SAB) in 5-fold serial dilutions on SAB agar and incubated at 30°C for 48 h to assess viability. CFUs before and after heat-ramp assays were enumerated to calculate relative survival. To determine growth differences at high temperature, untreated strains were spotted on SAB agar and incubated at 37°C for 48 h. Statistical significance was determined with the Anova test, via GraphPad Prism.

### RNA-seq analysis

RNA-seq datasets were downloaded from the GSEO database from the following deposited datasets: GSE162851, GSE136879, GSE93005, PRJEB4169, GSE32049, GSE32228, GSE121183, GSE60398, and GSE66510. Where necessary, raw data was reanalyzed by bowtie2 (2.3.5) (27) alignment to the most recent *C. neoformans* H99 or KN99α genome (fungibd.org), count matrices generated with HTSeq (1.99.2)(28), and RNA-seq analysis with Bioconductor DESeq2 (1.22.2) (29).

### Protein structure prediction and glycosylation analysis

Protein structure predictions were generated with AlphaFold2 v2.1.0 (reduced BFD database) (30). Intrinsically disordered regions were identified with IUPred (31), and glycosylation predictions were made with GPP (32)

## Data and code availability

Genome data can be accessed via accession PRJNA783275. This work did not lead to the generation of any new code.

## Results

The relationship of the original strains, previously described (7), is shown in **Figure 1**, with the original patient strain referred to as patient unpassaged (PU), and the original cockatoo strain referred to as cockatoo unpassaged (CU). Isolates recovered from mice infected with the CU strain are identified as cockatoo passaged brain (CPB) and cockatoo passaged lung (CPL) to denote the tissues from which they were isolated. *C. neoformans* recovered from patient and cockatoo are referred to as strains to denote different origins while those recovered from mouse tissues are referred to as isolates. Here, we aimed to identify the relatedness of the CU and PU strains, examine genetic changes resulting from mammalian passage of these strains, and interrogate differences across virulence phenotypes arising from passage across hosts and body sites.

**Figure 1.**
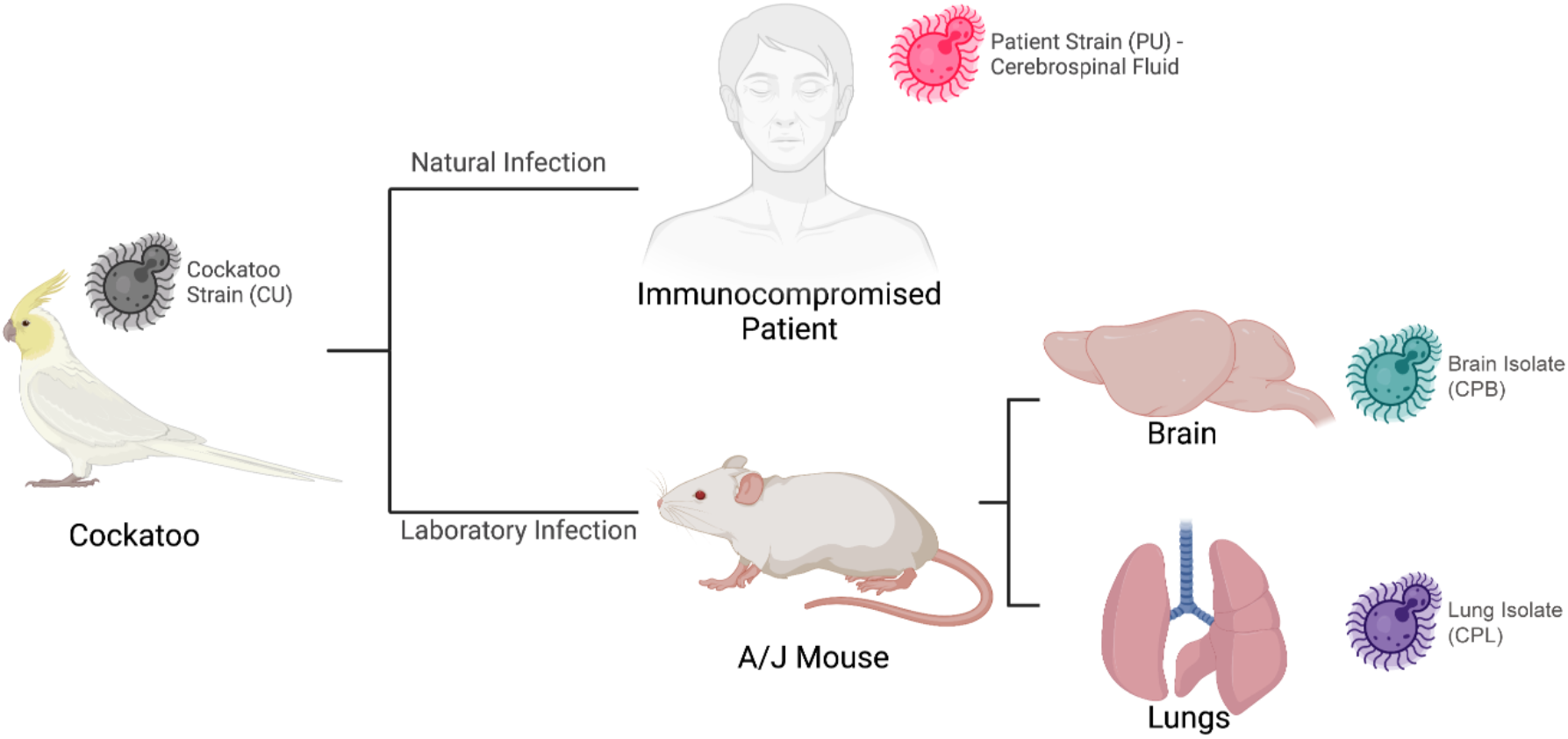
Passaging scheme and relationship between the *C. neoformans* strains and isolates. The cockatoo and patient strains had been kept frozen since the prior study was completed in 2000 (7). These are distinguished in that they were not passaged and are labelled cockatoo (CU) and patient (PU) where the U stands for unpassaged. The mouse passaged isolates come from a laboratory infection and are labelled CPB and CPL for cockatoo passaged brain and cockatoo passaged lung, respectively, to denote the mouse organ from where they were recovered.

### Genomic analysis of patient and cockatoo strains

To determine the relatedness of the PU and CU strains, Illumina reads were aligned to the *C. neoformans* H99 reference genome (33), and variants were called to identify single nucleotide polymorphisms (SNPs) and insertion/deletion events (indels). Based on a phylogenetic analysis of these samples and a subset of 238 additional whole genome sequences chosen to represent lineages VNIa, VNIb, VNIc, VNBI, VNBII, and VNII, previously described by Desjardins et al. (34), we assigned the PU and CU strains to the globally detected VNII lineage confirming their close relationship (**Fig. 2, Supplementary Fig. 1**), and determined that both strains possess the *MAT*α mating type. Utilizing ONT read data, we then generated genome assemblies for both PU and CU strains, consisting of complete telomere-to-telomere sequences for each chromosome, polished with Illumina reads. To identify variants, we compared the two assemblies with nucmer (MUMmer). We identified 7 genes with nonsynonymous variants between the two assemblies, all impacting genes with homologs in the *C. neoformans var. grubii* (H99) genome (**Table 1**). Of these genes, 2 (LQVO5_002184 and LQVO5_000317) had *C. neoformans* H99 homologs (CNAG_06273 and CNAG_00342) that are repressed during titan cell formation and murine cryptococcal infection, respectively (35, 36). For both strains, we were able to generate full-length chromosomal assemblies, consisting of 14 chromosomes, with rRNA content residing on chromosome 4, and chromosome 15 representing the mitochondrion, with equivalent gene sets and high levels of identity between the two assemblies (99.99%), indicating high levels of relatedness (**Supplementary Fig. 2a**). When these assemblies are compared to the *C. neoformans* H99 reference assembly, we see sequence identities of 97.7%, for both CU and PU strains to the H99 reference (**Supplementary Fig. 2b**).

### Mouse passage of cockatoo (CU) strain

The finding of genomic changes when comparing CU and PU strains suggested that the PU differences may have occurred during human infection. Consequently, we wondered whether passage of the CU strain in mice would result in similar genomic changes. A female A/J mouse was infected intravenously with the CU strain and observed for signs of illness, but none were apparent. Hence, at day 43 the mouse was sacrificed, brain and lung were harvested and homogenized, and the suspensions plated, which yielded 816 and 78 CFU/mg in brain and lungs, respectively. Three individual colonies were selected from the brain (CPB1-3) and lung (CPL1-3) CFU plates. These isolates were sequenced and studied for phenotypic characteristics.

**Table 1.** Genes with nonsynonymous variants between patient (PU) and cockatoo (CU) strains.

| CU(VNII) Gene | H99 (VNI) Homolog | Variant (CU>PU) | Function |
| --- | --- | --- | --- |
| **LQVO5_000317** | CNAG_00342 | 270C>G | KOG: SWI-SNF chromatin-remodeling complex protein |
| **LQVO5_001277** | CNAG_05000 | 1835T>C | EC: Exo-alpha-sialidase |
| **LQVO5_004692** | CNAG_05713 | 194C>G | RNA Polymerase II elongator complex protein 1 |
| **LQVO5_005253** | CNAG_02201 | 448C>G | KOG: Serine/threonine protein kinase |
| **LQVO5_005443** | CNAG_03102 | 122C>T | Hypothetical Protein |
| **LQVO5_006755** | CNAG_01999 | 34C>T | KOG: Alpha SNAP protein, membrane fusion |
| **LQVO5_002184** | CNAG_06273 | 217C>A<br>223C>T | Pfam: Ribosome biogenesis regulatory protein (RRS1) |

**Figure 2.**
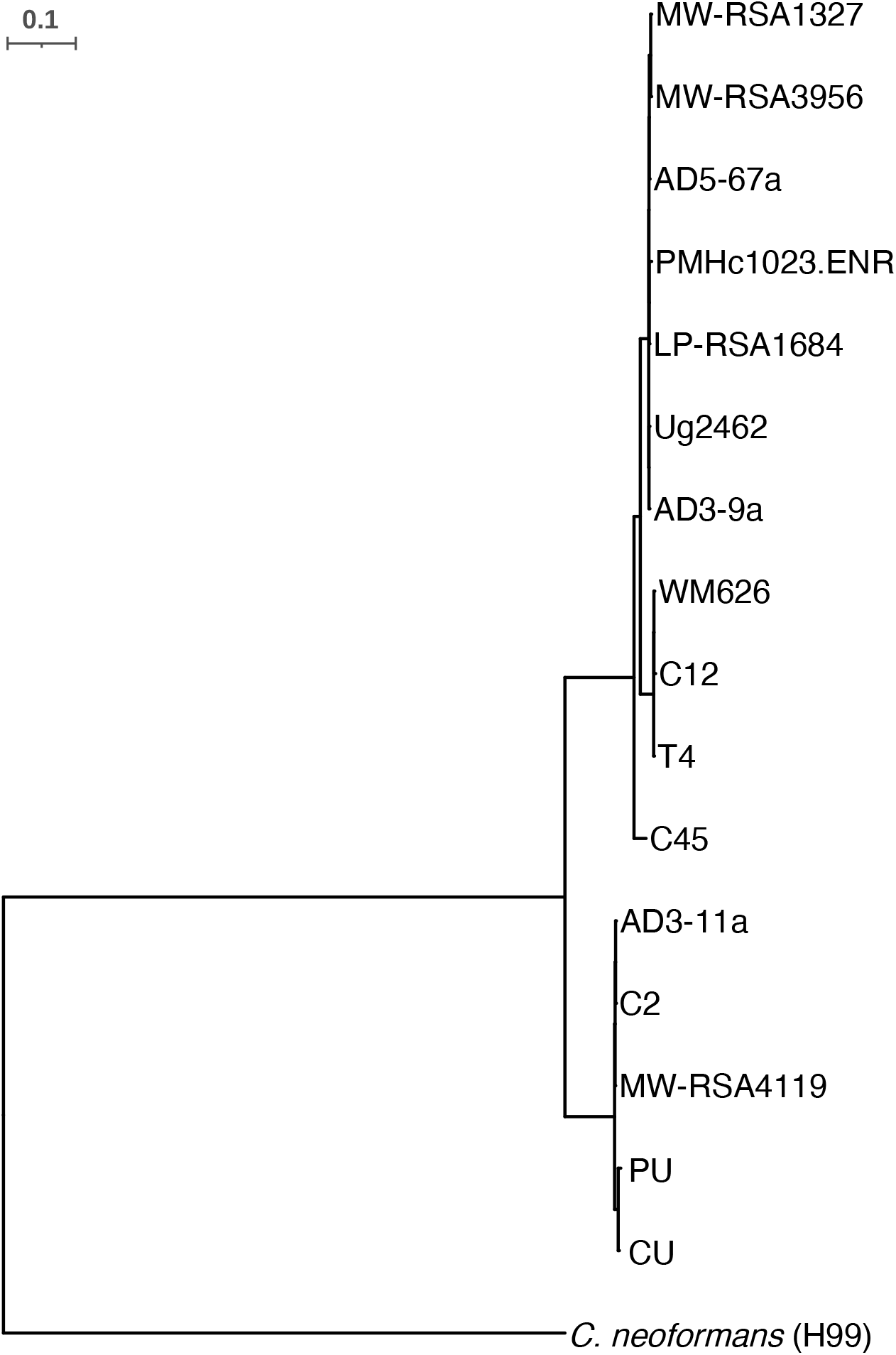
VNII phylogeny. SNP based maximum likelihood phylogeny for PU, CU and VNII strains rooted to the *C. neoformans* reference strain H99. Illumina reads for the VNII strains shown were aligned to the *C. neoformans* H99 reference genome to identify SNPs across the genome, which were used to infer a phylogeny with RAxML.

### Genomic analysis of mouse-passaged isolates

To identify variants arising from the passage of the CU strain in mice, we aligned Illumina data generated for all mouse evolved isolates to our CU genome assembly. Samples aligned with an average coverage of 775X across the CU reference. We found a frameshift variant in one of the seven genes altered in the patient strain, LQVO5_000317 (**Table 1**). This frameshift in LQVO5_000317 was present in all mouse evolved isolates from both brain and lung tissue (CPB1-3, CPL 1-3) (**Table 2**). The commonality of this variant across brain and lung isolates suggests the mutation was acquired at a common site prior to dissemination. This gene is a homolog of CNAG_00342 in the *C. neoformans* H99 (VNI) genome, which is predicted via eukaryotic orthologous groupings to function as a SWI-SNF chromatin-remodeling complex protein (KOG2510), and is down-regulated in a murine lung infection model (36), consistent with loss-of-function mutations in mammals.

**Table 2.**
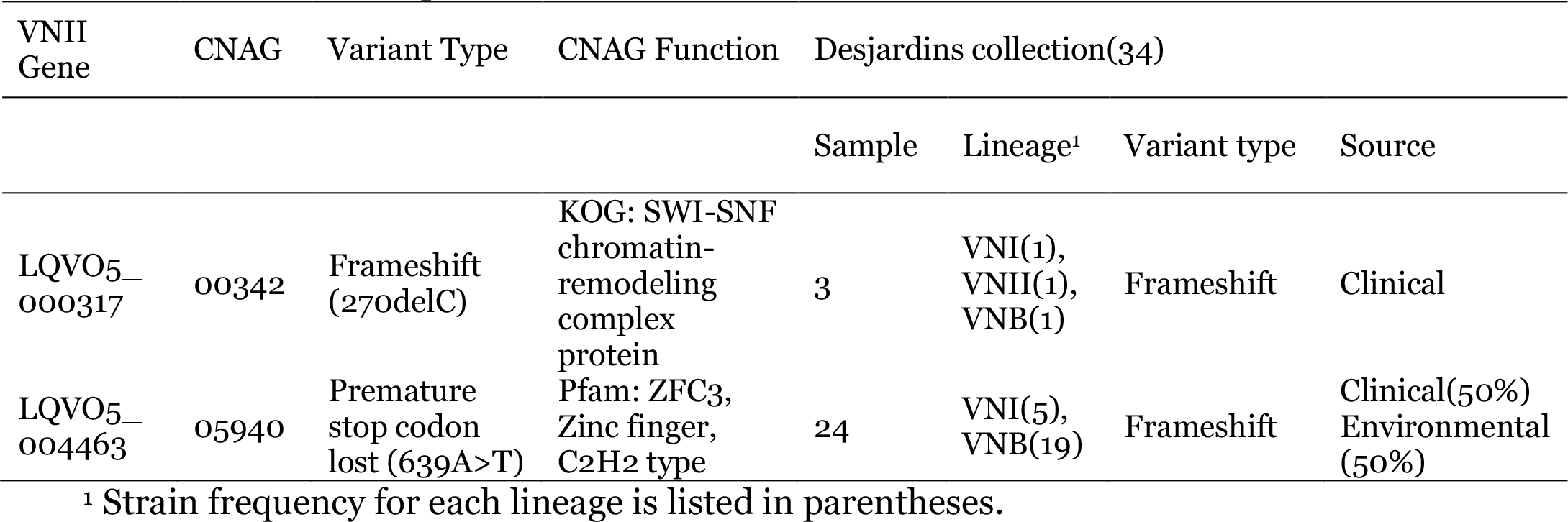
Variants in patient strain (PU) and mouse evolved isolates (CPB, CPL).

| VNII Gene | CNAG | Variant Type | CNAG Function | Desjardins collection(34) |  |  |  |
| --- | --- | --- | --- | --- | --- | --- | --- |
|  |  |  |  | Sample | Lineage<sup>1</sup> | Variant type | Source |
| LQVO5_000317 | 00342 | Frameshift (270delC) | KOG: SWI-SNF chromatin-remodeling complex protein | 3 | VNI(1), VNII(1), VNB(1) | Frameshift | Clinical |
| LQVO5_004463 | 05940 | Premature stop codon lost (639A>T) | Pfam: ZFC3, Zinc finger, C2H2 type | 24 | VNI(5), VNB(19) | Frameshift | Clinical(50%)<br>Environmental (50%) |

<sup>1</sup> Strain frequency for each lineage is listed in parentheses.

A second variant impacting only the mouse passaged isolates collected from lung tissue resulted in the loss of a premature stop codon in LQVO5_004463, a truncated homolog of CNAG_05940, a predicted Zinc-finger domain protein (ZFC3). A premature stop codon truncating the ZFC3 homolog (LQVO5_004463) is present in both the CU and PU strains, as well as the CPB isolates, but appears to have reverted to wild type in the CPL isolates. When LQVO5_004463 is extended through the loss of this premature stop codon, the resulting gene encodes a full-length protein comparable to CNAG_05940 in sequence and structure (**Supplementary Fig. 3**). Zfc3 has a single predicted zinc finger and a striking number of serine and threonine residues in the LQVO5_004463 homolog, CNAG_05940, totaling 21.9% of all residues present. The predicted structure of this protein includes long intrinsically disordered regions that may transition to structured regions upon binding to a substrate (**Supplementary Fig. 3c**). Interestingly, deletion of CNAG_05940 in *C. neoformans* results in strains with significantly increased capsule content (37), and this gene is thought to be a target of the virulence implicated transcription factors Gat201 and Liv3 (38). The presence of variants impacting shared genes in both patient and cockatoo-derived mouse isolates is consistent with the hypothesis that the patient’s *C. neoformans* infection was also derived from the pet cockatoo.

To assess the frequency of loss-of-function mutations in the SWI-SNF and ZFC3 homologs among clinical and environmental *Cryptococcus* isolates, we looked for loss-of-function variants in 387 published isolates from both patient and environmental sources (34). We found 3 clinical samples (from the lineages VNI, VNII, and VNB) with frameshift variants in CNAG_00342 (SWI-SNF) and 24 samples with frameshift variants present in CNAG_05940 (ZFC3) (**Table 2**). VNI and VNB isolates from both clinical and environmental sources are impacted by frameshift variants in CNAG_05940. To identify large-scale genomic variation in these CU, PU, CPB and CPL isolates, we looked for evidence of aneuploidy and copy number variation (CNV) based on sequence coverage, however, we saw no evidence of either aneuploidy or significant CNVs arising in response to human, bird, or murine passage.

### CNAG_00342 and CNAG_05940 expression in published datasets

To further probe the role of these genes altered in the patient and mouse isolates, we analyzed eight publicly deposited RNA-seq datasets of *C. neoformans* strains H99 and KN99α (**Table 3**). These databases were selected because they represented transcriptional studies done in conditions that resemble the expected conditions during infection. We found that both CNAG_00342 and CNAG_05940 were significantly differentially regulated (p < 0.05) under conditions related to those expected during animal passage and infection, namely 37 °C, increased CO_2_, in vitro infection of macrophages, or in vivo infection of rabbit CSF. Both genes were mostly upregulated in H99 strains and downregulated in KN99α rather than between conditions with the caveat that many of the H99 strains were exposed to ambient CO_2_ levels while both KN99α datasets were exposed to 5% CO_2_, indicating strain and condition specific changes that highlight variability in expression.

**Table 3.** Metadata from RNAseq datasets, only including Log2FC values with *P* < 0.05.

| Reference<br>PMID<sup>1</sup> | Strain | Conditions | Duration | CNAG_00342<br>Log<sub>2</sub>FC | CNAG_05940<br>Log<sub>2</sub>FC |
| --- | --- | --- | --- | --- | --- |
| 31666517 | H99 | 37 °C, YPD | 1 h | - | - |
| 28376087 | H99 | 37 °C, YPD | 2 h | 0.91 | - |
| 24743168 | H99 | 37 °C, YPD | Grown at<br>37 °C until<br>stationary | - | - |
| 31860441_CSF | H99 | Rabbit CSF | 24 h | 2.76 | 2.50 |
| 31860441 | H99 | 37 °C, 5%,<br>CO<sub>2</sub>,<br>Macrophage | 24 h | 6.30 | 4.40 |
| 22174677 | H99 | 37 °C, 5%<br>CO<sub>2</sub>, DMEM | 1.5 h | - | - |
| 27094327 | KN99α | 37 °C, 5%<br>CO<sub>2</sub>, DMEM | 1.5 h | - | -1.14 |
| 27094327_TM | KN99α | 37 °C, 5%<br>CO<sub>2</sub>, DMEM | 1.5, 3, 8,<br>24 h | 0.37 | -0.57 |
| 33688010 | KN99α | 37 °C, 5%<br>CO<sub>2</sub>, DMEM | 24 h | -4.75 | -3.23 |

<sup>1</sup>Dataset 31860441\_CSF indicates analysis of *C. neoformans* samples from cerebrospinal fluid *in vivo* infections, opposed to cryptococcal samples incubated in the described conditions *in vitro*. Dataset 27094327\_TM indicates analysis of an array of timepoints as a time course model, opposed to analysis of a single time point closest to the other datasets for consistency.

### Heat tolerance

To determine if resistance to cell death may be related to virulence in these strains, we compared the mouse passaged isolates to the patient strain in a cell death assay that has been previously demonstrated to induce gene-dependent cell death in *S. cerevisiae*, and more recently in *C. neoformans* (39, 40). To assess cell death susceptibility, a transient sublethal heat-ramp (not heat shock) was applied to all isolates and survival was determined by CFUs when plated at 30°C. Interestingly, all mouse, cockatoo, and patient strains were death-resistant when compared to the lab strain, KN99α (**Fig. 3A**). Among the isolates tested, the three mouse-passaged brain isolates (CPB1-3) were significantly more death-resistant than the lung isolates (CPL1-3), and the patient (PU) strain (p=0.0324 and p = 0.002, test=ANOVA) (**Fig. 3B**). However, cell death-resistance does not appear to reflect a gain of heat tolerance at body temperature as all isolates grew indistinguishably when untreated samples were plated on SAB agar and incubated at 37°C (**Fig. 3C**). Although the lab strain KN99α may be slightly more robust than the CU, PU, CPB, and CPL isolates at both 30°C and 37°C, no differences in CFU number or size among the isolates was observed, suggesting that heat-ramp cell death resistance is not directly correlated with the ability to grow at high temperature.

**Figure 3.**
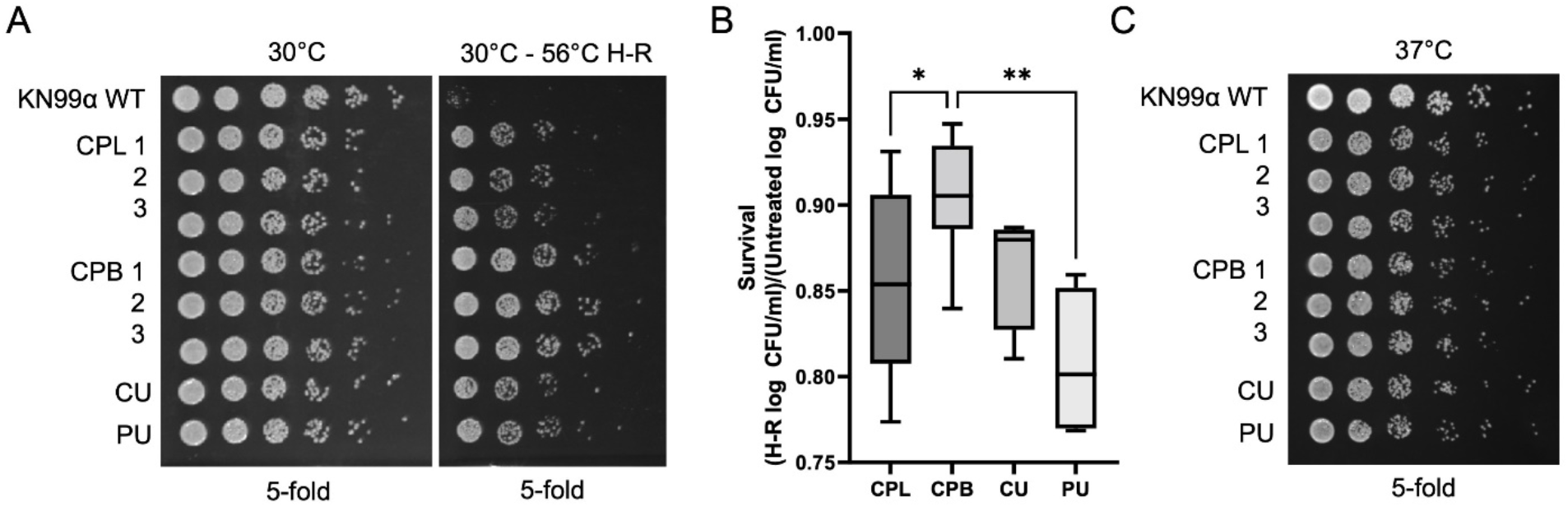
Heat ramp experiments to evaluate thermal survival of the various *C. neoformans* strains and isolates. The cockatoo passage brain isolates (CPB) recovered from mice infected with the cockatoo strain (CU) were more resistant to cell death when compared to lung isolates (CPL) or to the original cockatoo (CU) and patient strains (PU). (A) Survival after a cell death stimulus [10 min linear 30° to 56°C heat-ramp (H-R)] was determined by CFU on SAB agar plates incubated 2 days at 30°C. (B) Quantification for A calculated as the ratio of log 10 CFU/ml of heat-ramp treated to untreated from four independent experiments. Ordinary one-way ANOVA with Tukey’s multiple comparisons test, *p = 0.0139 **p = 0.0012. (C) Growth of the same isolates (no heat-ramp) spotted on SAB and incubated for 2 days at 37°C, representative of two independent experiments; no differences between isolates were detected.

### Growth in vitro and virulence factor expression

We analyzed both the CU and PU strains and the CPB and CPL isolates for growth in vitro (**Fig. 4**) and expression of four phenotypes known to be associated with virulence factors: capsule, melanin, urease, and phospholipase (**Figs. 5, 6**). All four isolates grew well in culture with the CPB1 isolate recovered from brain tissue growing faster than the others (**Fig. 4**). All isolates expressed each of these virulence factors but there were subtle differences observed. The original cockatoo strain (CU), patient strain (PU), and mouse passaged CU derivative isolates varied little with regards to the capsule, except that CPL1 cells had significantly smaller capsules than the other isolates (p=1×10^−4^, 4×10^−8^, 3.8×10^−6^, CPL1 vs CU, PU, CPB1, respectively; test= paired t-test) (**Fig. 5**), consistent with a functional ZFC3. Since a prior analysis of sequential isolates recovered from individual patients showed changes in polysaccharide structure (41), we analyzed their exopolysaccharide (EPS) by NMR, which revealed variation in O-acetylation content such that CU (37.46) > PU (26.15) > CPL1 (23.01) > CPB1 (19.50) (**Fig. 5A**). Further analysis of the Structural Reporter Group (SRG) ^1^H NMR resonances corresponding from backbone mannose anomeric protons, which provides a spectral signature for the repeating triad of glucuronoxylomannan (4.8-5.4ppm) (42) showed the same peak-set for CU and PU strains and CPB and CPL isolates (4.91, 4.97, 4.99, 5.10, 5.12, 5.15. 5.16, 5.20) (**Fig. 5C**), implying conservation of EPS structure in these strains and isolates. Isolates from the brain of the CU-infected mouse, CPB1-3, had faster rates and higher total percentages of melanization when compared to the other isolates, including the parental CU strain, the PU strain, and the CPL1-3 isolates from the lungs of the mouse (**Fig. 6A-C**). The CPB1 isolate was significantly more melanized than CU at 37°C when grown on L-DOPA-agar (p=0.0365, 0.0009, 0.0002; 2, 3, 6 d; test = ANOVA). For phospholipase, the size of the precipitation zone varied across isolates and increased with the time of incubation. After 72h of incubation at 30°C, the isolate recovered from the mouse brain (CPB1) presented significantly lower (p = 0.033, test=t-test) phospholipase activity than the original CU strain. After 96h the phospholipase activity of CPB1 was significantly lower (p = 0.0049, test=t-test) than in all other strains and isolates (**Fig. 6D**). In contrast, the expression of urease was comparable among the four isolates (**Fig. 6E**).

**Figure 4.**
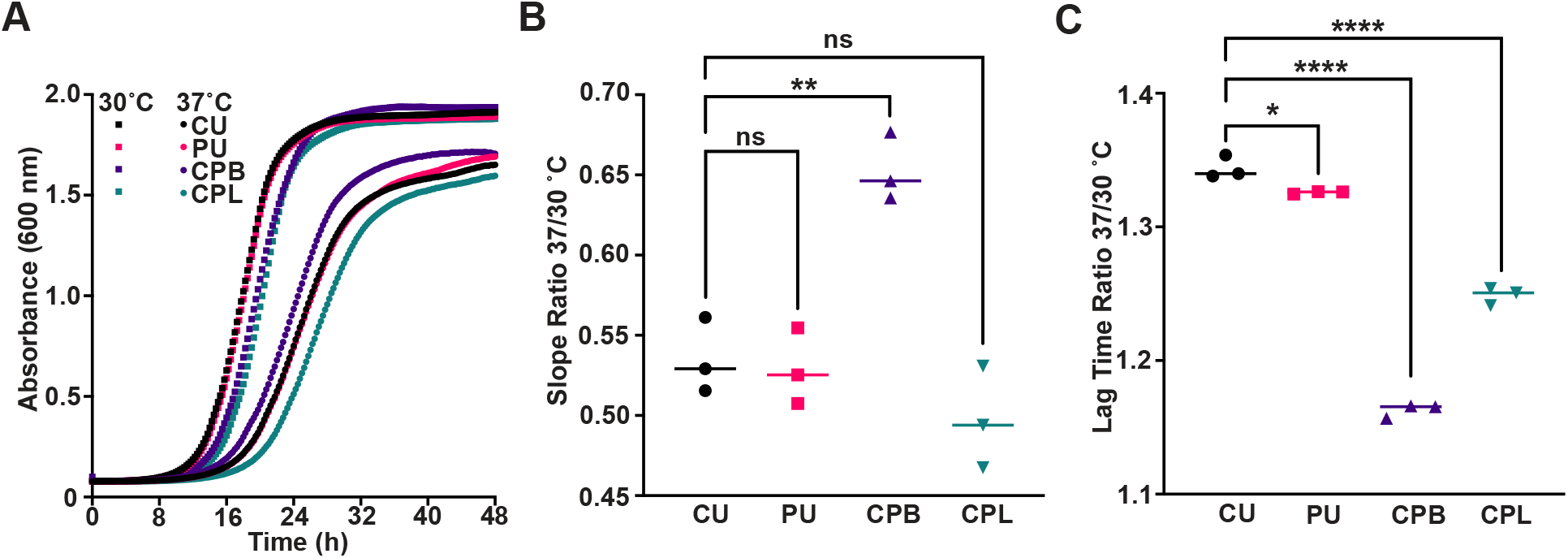
Comparison of *C. neoformans* cockatoo, human, and mouse-passaged derivatives growth at 30°C and 37°C. (A) Growth curves of *C. neoformans* strains cultured in Sabouraud dextrose broth at the indicated temperatures plotted as the mean of three biological replicates. (B) Comparison of growth rates at the two temperatures expressed as a ratio of linear phase slopes indicates a significant growth advantage at 37°C for the mouse-passaged brain isolate compared to the other strains and isolates. (C) Both the patient strain and mouse-derived isolates show a significant decrease in the length of lag time at 37°C compared to 30°C. Statistical significance was determined using an ordinary one-way ANOVA (ns = not significant, *p < 0.05, **p < 0.01, ***p < 0.001, ****p < 0.0001).

**Figure 5.**
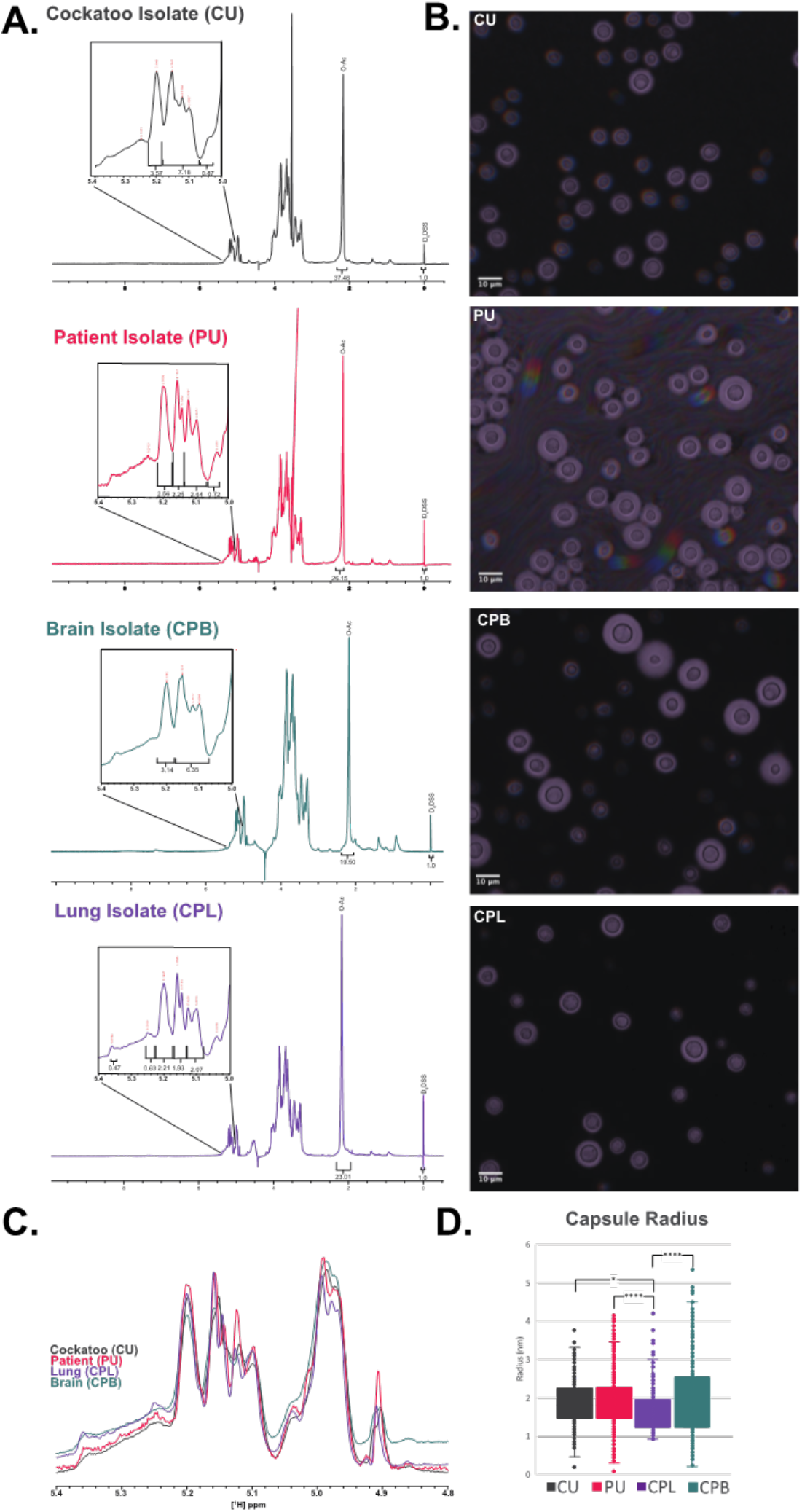
Capsular Characterization of Cockatoo and Patient strains, and Mouse passage isolates. (A) 1D [^1^H] NMR analysis of EPS isolated from CU and derivative isolates. Integration of peaks and comparison to internal standard (D_6_DSS) shows variation in polysaccharide O-acetylation. (B) India ink microscopy images of CU and derivative strains. (C) Overlay of structural reporter group (SRG) region of 1D [1H] NMR spectra indicates very little variation between strains with regards to GXM motifs.(D) Quantitative capsule analysis (QCA) indicates that the capsules of mouse passaged lung isolates are smaller than other cockatoo and derivative isolates.

**Figure 6.**
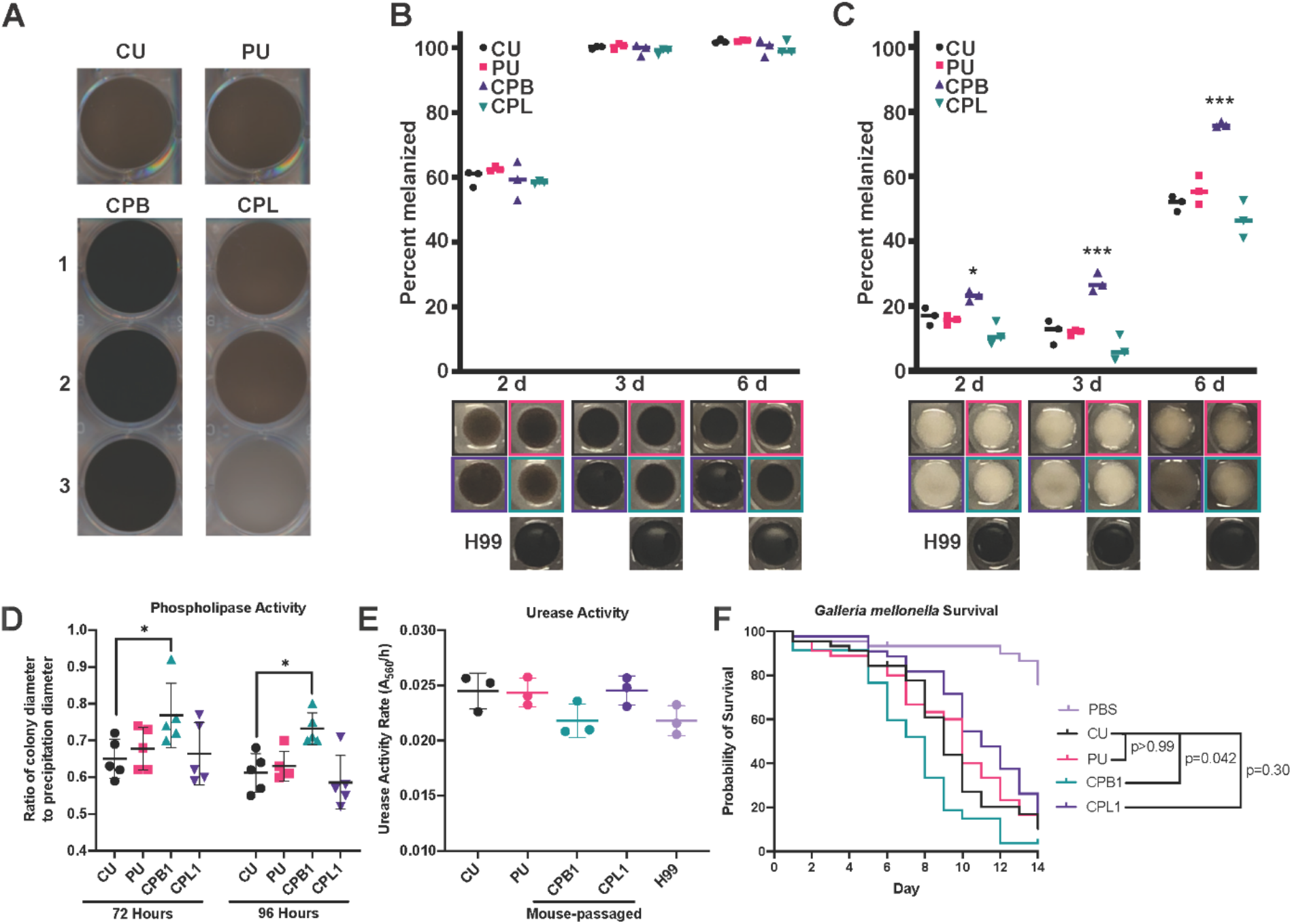
Changes in virulence factor activity associated with passaging the cockatoo strain in mice. (A) Melanization is greatly enhanced in the CPB isolates after growth at 30°C for 5 days. There are no major differences in melanization between the PU strain or CPL isolates, which generally show melanization consistent with the parental strain (CU). (B-C) Scatter plot graphs (upper panel) and representative images (lower panels) of pigment production expressed as a percentage of the H99 reference strain for the cockatoo (CU) and human (PU) strains, and mouse-passaged brain (CPB) and lung (CPB) isolates grown for the indicated amount of time on L-DOPA-agar at either 30°C (B) or 37°C (C). Compared to the original cockatoo strain, only the mouse-derived brain isolate shows a significant increase in pigmentation at 37°C. (D) *C. neoformans* were inoculated onto egg yolk agar and incubated at 30C. After 72h and 96h of incubation phospholipase production was analyzed by measuring the ratio of colony diameter to precipitate + colony diameter on the plate, where a ratio value equal to 1.0 indicates a lack of phospholipase activity. (E) Time course of urease activity for the indicated strains of *C. neoformans* grown at 30°C in urea broth. Increased pH of culture media that results from the conversion of urea to ammonium was quantified by measuring the absorbance of cell culture media at 560 nm relative to a cell-free control. Urease activity rates were not statistically different between strains. (F) There is no statistical difference in virulence between the CU and PU strains in the *G. mellonella* model system. However, the CPB1 isolate from the mouse had significantly enhanced virulence when compared to the parental CU strain. Log-rank Mantel-Cox test was performed using GraphPad PRISM and corrected for multiple comparisons using the Bonferroni method.

### Virulence in *Galleria mellonella*

To identify strain differences in virulence, we tested the CU, PU, CPB, and CPL isolates in the invertebrate model *G. mellonella* (**Fig. 6F**). This model has been used previously to compare virulence of *Cryptococcus* isolates, which roughly correlates to virulence in mammalian models (1,2). We found no statistical differences between the virulence of the CU and PU strains. There was enhanced virulence of the CPB1 brain isolate compared to its parental CU strain, and reduced virulence of the CPL1 lung isolate. While all strains and isolates tested were virulent, these results show subtle changes in virulence for an insect host.

### Interactions with macrophages

The interaction of *C. neoformans* with macrophages is unusual in that ingestion results in transient intracellular residence, which may be followed by non-lytic exocytosis whereby the fungal cell exits the phagocytic cell without lysing the latter (43, 44). Given that this process involves a complex choreography of cellular events that must occur in synchrony we considered it a sensitive indicator of *C. neoformans*-macrophage interaction, and examined its frequency for the CU, PU, CPL, and CPB isolates (**Fig. 7**). The results show that the CPB1 isolate manifested a significantly higher frequency of non-lytic exocytosis relative to the other strains and isolates (p=0.013, test of equal proportions with Bonferroni correction).

**Figure 7.**
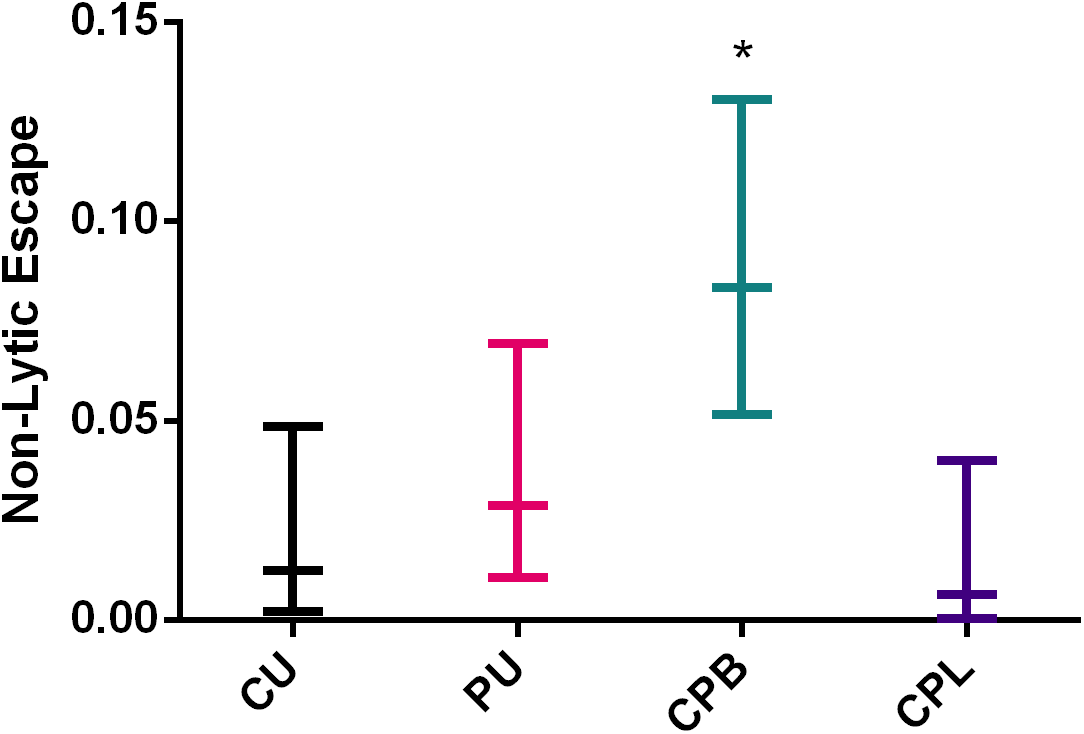
Non-Lytic escape frequency of each strain during BMDM infection. CPB was the only isolate with a significantly increased frequency of non-lytic escape compared to the original CU strain. * Indicates *P* < 0.05 via a test of equal proportions with Bonferroni correction.

### Interactions with amoeba

Since amoebae are predators of *C. neoformans* and amoeba-fungal interactions have been proposed to select for traits that function during mammalian virulence, we evaluated whether passage through human and mice affected the interaction with *A. castellanii*. In an assay favoring amoeba predation through the presence of divalent cations (45), we observed that *C. neoformans* strains recovered from the patient (PU) and cockatoo (CU), as well as mouse-passaged isolates (CPL and CPB) were equally susceptible to predation (**Fig. S4**). All fungal strains experienced minimal killing by *A. castellanii* at 24 h, and modest CFU rebounds by 48 h, but differences between the CU and PU strains or CPL and CPB isolates were not significant (**Fig. S4A**). This pattern of initial CFU decrease and subsequent rebound was not seen for the same strains or isolates incubated in only DPBS (**Fig. S4B**).

## Discussion

Comparison of the patient and cockatoo strains revealed that they are genetically and phenotypically very similar and likely derived from each other. Given that the patient strain came from an immunocompromised patient with cryptococcosis, and the cockatoo strain came from bird excreta in the home of the patient, the inference was made that the bird was the source of the infection (7). Although psittacine birds can develop cryptococcosis, their association with *C. neoformans* is usually in the form of saprophytic growth in their excrement (46). In this situation, the occurrence of cryptococcosis in an immunocompromised individual who lived in close proximity to the cockatoo was inferred to be a case of human infection from exposure to a point source associated with the pet bird (7). The near genomic identity of the patient and cockatoo strains confirms the finding that they were indistinguishable based on restriction length polymorphisms and supports this scenario (7). However, the prior case report could not rule out the possibility that the patient had acquired the infection first and then infected the bird in some form, possibly through cough during a pulmonary phase of disease. Although this scenario was, and is, considered unlikely, the observation that mice with experimental cryptococcosis contaminate their cage bedding indicates that hosts with systemic infection can shed *C. neoformans* (47), providing some support for the plausibility of patient to bird transmission. However, our finding that some of the mutations observed when the cockatoo strain was passaged in mice are also found in the patient strain suggests that these were selected in mammalian hosts, which in turn supports the original conclusion that the transmission was from the bird to the patient.

The comparison of patient and cockatoo strains revealed amino acid changes in 7 proteins. There were two non-synonymous changes in the predicted ribosome regulatory protein (LQVO5_002184, homolog of the H99 gene CNAG_06273), a gene repressed during titan cell formation (35). One variant is found in the predicted SWI-SNF chromatin-remodeling complex (LQVO5_000317, homolog of the H99 gene CNAG_00342), a protein that is down-regulated during murine cryptococcal infection (36), is involved in cryptococcal morphogenesis (48), and is implicated in nitrosative stress response in the plant pathogenic fungus *Fusarium graminearum* (49).

To ascertain whether we could reproduce some of the changes observed in the patient strain relative to the cockatoo strain we passaged the latter in mice. In all mouse passaged isolates we observed one frameshift variant that impacts the same gene (LQVO5_000317, homolog of the H99 gene CNAG_00342) observed in the patient strain. A second variant impacting only the mouse passaged isolates collected from lung tissue results in the loss of a premature stop codon for LQVO5_004463, a homolog of CNAG_05940 (ZFC3). ZFC3 is a transcription factor that is repressed during titan cell induction in addition to being a target of the transcription factors Gat201 and Liv3 (38). Changes in length of the protein encoded by LQVO5_000317 are likely to impact expression of this gene’s targets. Analysis of the expression of CNAG_00342 and CNAG_05940 in publicly available databases involving conditions related to animal passage and infection revealed variation in the expression of these genes, however, it is unclear whether the condition or genetic background is responsible for this variation. Hence, it may be that the genomic changes observed in the cockatoo strain upon mouse or human passage represent selection of this particular genotype during infection, which may be specific to this genetic background. Variation of gene expression across strains, correlating with genetic groupings and lineage, have been observed for clinical and environmental isolates (50), and isolates derived directly from human CSF (51). Consequently, we caution against generalization from the results to other cryptococcal strains until there is greater sampling of changes associated with virulence; given the plasticity of the *C. neoformans* genome, there may be many solutions to the problem of survival in the ecologic niche defined by these hosts.

The occurrence of numerous *C. neoformans* genetic changes in the form of SNPs, deletions, and insertions suggests that fungal replication in mammalian hosts may select for specific changes. Although this is the first genomic study of a *C. neoformans* strain before and after human passage, other studies have reported genetic changes during infection. Analysis of serial isolates from patients shows chromosome rearrangements, ploidy alterations, SNPs, insertions, and deletions (52). One mechanism of mutagenesis is transposon mobilization during infection (53), however, we saw no evidence of transposon mobilization upon mammalian passage. *C. neoformans* replicates within macrophages in vivo (54), and internalized fungal cells are exposed to oxygen-and nitrogen-derived radicals that can be mutagenic (55). A recent study of *C. albicans* during in vitro and in vivo passage suggested a higher mutation rate in vivo (56). The fact that mammalian infection may select for specific genetic changes in *C. neoformans* suggests that those microbes capable of prolonged residence in hostile environments such as human hosts can acquire genetic changes, such that the capacity for virulence could provide a shortcut to greater genetic variation.

Our observation that human and mouse passage was associated with the emergence of genetically different variants has important implications for the understanding of *C. neoformans* genomics, virulence, and pathogenesis. Furthermore, the finding that passage of an environmental strain of *C. neoformans* through humans and mice resulted in genetic changes suggests that clinical strains may have been modified by residence in human hosts, where they are exposed to higher constant temperature and must survive attack by the immune system. However, these changes did not confer increased resistance to amoeba, in contrast with previous findings that the passage of environmental isolates in amoebae results in both genetic changes and the emergence of pleiotropic phenotypes (57). Since *Entamoeba* spp. and slime molds can occur in bird feces (58, 59), the CU strain may already be maximally resistant to amoeba predation. Comparison of genomic changes in mammalian and amoeba passaged *Cryptococcus* revealed no obvious commonalities, consistent with the notion that even whilst these hosts provide similar challenges to *C. neoformans*, such as surviving phagocytosis and phagosome residence, they constitute different selective environments. Overall, these results indicate that the cryptococcal genome is highly malleable such that genetic changes can accumulate rapidly.

Comparison of clinical and environmental isolates for genomic differences such as the ones found in this study revealed multiple instances of frameshift variants in both CNAG_00342 and CNAG_05940 in clinical and environmental isolates spanning lineages VNI, VNII, and VNB (34). How these genetic changes affect virulence and pathogenesis is a question for future studies. In this regard, a comparison of the virulence of 10 clinical and 11 environmental *C. neoformans* isolates in mice revealed that 7 clinical isolates and only 1 environmental isolate caused lethal infection (60). Considering these results in the light of our findings suggest that the clinical strains in that study were perhaps more virulent because of genetic changes that occurred or were selected for by human passage.

The comparison of *C. neoformans* characteristics associated with virulence among cockatoo, patient, and mouse passaged isolates in this study revealed subtle phenotypic changes. There were differences in average capsule size between isolates recovered from mouse lung and brain, but NMR analysis of the major polysaccharide component revealed no major changes. The finding of organ related differences in capsule size is consistent with prior reports (61). However, the finding that the GXM structure was unchanged contrasted with the prior observation that serial clinical isolates from patients with persistent infection manifested changes in the polysaccharide structure (41), suggesting that for the CU strain studied here the polysaccharide structure was more stable. The major difference in the polysaccharide structures involved the extent of acetylation, with the CPB isolate having more than others. No major differences were observed in urease expression, but the mouse-passaged brain isolates manifested faster melanization. Despite melanin variability across the CPL isolates, we did not see genetic variation within genes known to be involved in melanization in these isolates. Similarly, mouse-passaged isolates and patient strains grew better at higher temperatures than the cockatoo strain possibly reflecting a period of adaptation to thermal mammalian conditions. In general, there were no major changes in phenotypes associated with virulence during mammalian passage consistent with the notion that these attributes exist primarily for environmental survival and only accidentally confer upon *C. neoformans* the capacity for mammalian virulence. Nevertheless, we do note that the mouse passaged isolates recovered from brain tissue manifested faster growth, higher melanization, increased rates of non-lytic escape, and killed moth larvae faster, consistent with a relative gain in virulence during animal passage. These inter-strain and -isolate phenotypic differences highlight the tremendous variation apparent in closely related *C. neoformans* strains, a phenomenon that contributes to virulence (62) and is also apparent in pleiotropic variants generated by amoeba predation (57) and phenotypic switching (63).

We note with interest that the mouse passaged isolates recovered from brain tissue (CPB) were more resistant to thermal stress. Since the mouse passaged isolates recovered from lung tissue (CPL) did not show this phenotype we cannot attribute this to simple thermal adaptation to mammalian temperatures. Furthermore, the comparison of genetic variants between CPB and CPL isolates did not reveal genetic changes that are known to confer increased thermal stability. Consequently, the most likely explanation for increased thermal tolerance in CPB isolates is epigenetic change, possibly associated with altered metabolic states, which may allow for greater survival during rapid heating. Brain and lung environments are expected to differ in catecholamine concentrations (64) and inflammatory responses (65-67). Although a mechanistic investigation of this phenomenon is beyond the scope of this paper, we note that if this phenomenon occurs in nature, it could provide environmental fungi capable of mammalian infection with a mechanism for rapid thermal adaptation that could increase their fitness during climate change. This in turn raises the specter that fungi capable of mammalian infection could increase in prevalence with warming climates.

In summary, genomic analysis of cockatoo, human, and mouse passaged isolates strongly supports our earlier inference that human infection resulted from exposure to a pet cockatoo. In this study, the comparison of cryptococcal genomes of the incident bird and patient strains with mouse-passaged isolates revealed the occurrence of common genetic changes during mammalian passage. Previously we showed that passage of *C. neoformans* in mice promotes the appearance of new electrophoretic karyotypes (69). Similarly, in the ascomycete *C. albicans*, infection was associated with larger changes in heterozygosity during murine passage (56) and human infection (70) than occur in vitro. The capacity for virulence in pathogenic microbial species is not without cost as evident by genome reduction and host specialization (71). The finding that mammalian infection promotes genomic changes in both *C. neoformans* and *C. albicans* suggests that the capacity for virulence can provide a mechanism for more rapid evolutionary change through selection and adaptation in the mammalian host, which brings new parameters for consideration when evaluating the cost-benefit equation for mammalian virulence in pathogenic fungi.

## References

1. Casadevall A, and Perfect JR. Cryptococcus neoformans. Washington, DC: American Society for Microbiology; 1998.

2. Goldman DL, Khine H, Abadi J, Lindenberg DJ, Pirofski L, Niang R, and Casadevall A. Serologic evidence for Cryptococcus infection in early childhood. Pediatrics. 2001;107(E66.

3. Alanio A. Fungal latency. JClinInvest. 2020;in press(

4. Currie BP, Freundlich LF, and Casadevall A. Restriction fragment length polymorphism analysis of Cryptococcus neoformans isolates from environmental (pigeon excreta) and clinical isolates in New York City. J Clin Microbiol. 1994;32(1188–92.

5. Alanio A. Dormancy in Cryptococcus neoformans: 60 years of accumulating evidence. The Journal of clinical investigation. 2020;130(7):3353–60.

6. Farrer RA, Borman AM, Inkster T, Fisher MC, Johnson EM, and Cuomo CA. Genomic epidemiology of a Cryptococcus neoformans case cluster in Glasgow, Scotland, 2018. Microbial genomics. 2021;7(3).

7. Nosanchuk JD, Shoham S, Fries BC, Shapiro DS, Levitz SM, and Casadevall A. Evidence for zoonotic transmission of Cryptococcus neoformans from a pet cockatoo to an immunocompromised patient. Ann Intern Med. 2000;132(205–8.

8. Littman ML, and Schneierson SS. Cryptococcus neoformans in pigeon excreta in New York City. Am J Hyg. 1959;69(49–59.

9. Kielstein P, Hotzel H, Schmalreck A, Khaschabi D, and Glawischnig W. Occurrence of Cryptococcus spp. in excreta of pigeons and pet birds. Mycoses. 2000;43(1-2):7–15.

10. Shrestha RK, Stoller JK, Honari G, Procop GW, and Gordon SM. Pneumonia due to Cryptococcus neoformans in a patient receiving infliximab: possible zoonotic transmission from a pet cockatiel. Respir Care. 2004;49(6):606–8.

11. Vanstraelen K, Lagrou K, Maertens J, Wauters J, Willems L, and Spriet I. The Eagle-like effect of echinocandins: what’s in a name? Expert review of anti-infective therapy. 2013;11(11):1179–91.

12. Cuomo CA, Rhodes J, and Desjardins CA. Advances in Cryptococcus genomics: insights into the evolution of pathogenesis. Memorias do Instituto Oswaldo Cruz. 2018;113(7):e170473.

13. Dragotakes Q, and Casadevall A. Automated Measurement of Cryptococcal Species Polysaccharide Capsule and Cell Body. Journal of visualized experiments : JoVE. 2018131).

14. Velegraki A, Kambouris M, Kostourou A, Chalevelakis G, and Legakis NJ. Rapid extraction of fungal DNA from clinical samples for PCR amplification. Med Mycol. 1999;37(1):69–73.

15. Fisher S, Barry A, Abreu J, Minie B, Nolan J, Delorey TM, Young G, Fennell TJ, Allen A, Ambrogio L, et al. A scalable, fully automated process for construction of sequence-ready human exome targeted capture libraries. Genome Biol. 2011;12(1):R1.

16. Koren S, Walenz BP, Berlin K, Miller JR, Bergman NH, and Phillippy AM. Canu: scalable and accurate long-read assembly via adaptive k-mer weighting and repeat separation. Genome Res. 2017;27(5):722–36.

17. Walker BJ, Abeel T, Shea T, Priest M, Abouelliel A, Sakthikumar S, Cuomo CA, Zeng Q, Wortman J, Young SK, et al. Pilon: an integrated tool for comprehensive microbial variant detection and genome assembly improvement. PLoS One. 2014;9(11):e112963.

18. Delcher AL, Phillippy A, Carlton J, and Salzberg SL. Fast algorithms for large-scale genome alignment and comparison. Nucleic Acids Res. 2002;30(11):2478–83.

19. Li H. Aligning sequence reads, clone sequences and assembly contigs with BWA-MEM. arXiv preprint 13033997. 2013.

20. Cingolani P, Platts A, Wang le L, Coon M, Nguyen T, Wang L, Land SJ, Lu X, and Ruden DM. A program for annotating and predicting the effects of single nucleotide polymorphisms, SnpEff: SNPs in the genome of Drosophila melanogaster strain w1118; iso-2; iso-3. Fly. 2012;6(2):80–92.

21. Stamatakis A. RAxML version 8: a tool for phylogenetic analysis and post-analysis of large phylogenies. Bioinformatics (Oxford, England). 2014;30(9):1312–3.

22. Yu G, Smith DK, Zhu H, Guan Y, and Lam TTY. ggtree: an R package for visualization and annotation of phylogenetic trees with their covariates and other associated data. Methods in Ecology and Evolution. 2017;8(1):28–36.

23. Abyzov A, Urban AE, Snyder M, and Gerstein M. CNVnator: an approach to discover, genotype, and characterize typical and atypical CNVs from family and population genome sequencing. Genome Res. 2011;21(6):974–84.

24. Roberts GD, Horstmeier CD, Land GA, and Foxworth JH. Rapid urea broth test for yeasts. Journal of clinical microbiology. 1978;7(6):584–8.

25. Chen SCA, Muller M, Zhou JZ, Wright L, and Sorrell TC. Phospholipase activity in Cryptococcus neoformans: a new virulence factor? J Infect Dis. 1997;175(414–20.

26. Malliaris SD, Steenbergen JN, and Casadevall A. Cryptococcus neoformans var. gattii can exploit Acanthamoeba castellanii for growth. Med Mycol. 2004;42(149–58.

27. Langmead B, and Salzberg SL. Fast gapped-read alignment with Bowtie 2. Nature methods. 2012;9(4):357–9.

28. Anders S, Pyl PT, and Huber W. HTSeq--a Python framework to work with high-throughput sequencing data. Bioinformatics (Oxford, England). 2015;31(2):166–9.

29. Love MI, Huber W, and Anders S. Moderated estimation of fold change and dispersion for RNA-seq data with DESeq2. Genome Biol. 2014;15(12):550.

30. Jumper J, Evans R, Pritzel A, Green T, Figurnov M, Ronneberger O, Tunyasuvunakool K, Bates R, Žídek A, Potapenko A, et al. Highly accurate protein structure prediction with AlphaFold. Nature. 2021;596(7873):583–9.

31. Mészáros B, Erdos G, and Dosztányi Z. IUPred2A: context-dependent prediction of protein disorder as a function of redox state and protein binding. Nucleic Acids Res. 2018;46(W1):W329–w37.

32. Hamby SE, and Hirst JD. Prediction of glycosylation sites using random forests. BMC bioinformatics. 2008;9(500.

33. Janbon G, Ormerod KL, Paulet D, Byrnes EJ, 3rd, Yadav V, Chatterjee G, Mullapudi N, Hon CC, Billmyre RB, Brunel F, et al. Analysis of the genome and transcriptome of Cryptococcus neoformans var. grubii reveals complex RNA expression and microevolution leading to virulence attenuation. PLoS genetics. 2014;10(4):e1004261.

34. Desjardins CA, Giamberardino C, Sykes SM, Yu CH, Tenor JL, Chen Y, Yang T, Jones AM, Sun S, Haverkamp MR, et al. Population genomics and the evolution of virulence in the fungal pathogen Cryptococcus neoformans. Genome Res. 2017;27(7):1207–19.

35. Trevijano-Contador N, de Oliveira HC, Garcia-Rodas R, Rossi SA, Llorente I, Zaballos A, Janbon G, Arino J, and Zaragoza O. Cryptococcus neoformans can form titan-like cells in vitro in response to multiple signals. PLoS Pathog. 2018;14(5):e1007007.

36. Li H, Li Y, Sun T, Du W, Li C, Suo C, Meng Y, Liang Q, Lan T, Zhong M, et al. Unveil the transcriptional landscape at the Cryptococcus-host axis in mice and nonhuman primates. PLoS neglected tropical diseases. 2019;13(7):e0007566.

37. Jung KW, Yang DH, Maeng S, Lee KT, So YS, Hong J, Choi J, Byun HJ, Kim H, Bang S, et al. Systematic functional profiling of transcription factor networks in Cryptococcus neoformans. Nature communications. 2015;6(6757.

38. Homer CM, Summers DK, Goranov AI, Clarke SC, Wiesner DL, Diedrich JK, Moresco JJ, Toffaletti D, Upadhya R, Caradonna I, et al. Intracellular Action of a Secreted Peptide Required for Fungal Virulence. Cell host & microbe. 2016;19(6):849–64.

39. Teng X, Cheng WC, Qi B, Yu TX, Ramachandran K, Boersma MD, Hattier T, Lehmann PV, Pineda FJ, and Hardwick JM. Gene-dependent cell death in yeast. Cell death & disease. 2011;2(8):e188.

40. Stolp ZD, Kulkarni M, Liu Y, Zhu C, Jalisi A, Lin S, Casadevall A, Cunningham KW, Pineda FJ, Teng X, et al. Gene-dependent yeast cell death pathway requires AP-3 vesicle trafficking leading to vacuole membrane permeabilization. bioRxiv. 2021:2021.08.02.454728.

41. Cherniak R, Morris LC, Belay T, Spitzer ED, and Casadevall A. Variation in the structure of glucuronoxylomannan in isolates from patients with recurrent cryptococcal meningitis. Infect Immun. 1995;63(1899–905.

42. Cherniak R, Valafar H, Morris LC, and Valafar F. Cryptococcus neoformans chemotyping by quantitative analysis of 1H NMR spectra of glucuronoxylomannans using a computer simulated artificial neural network. Clin Diagn Lab Immunol. 1998;5(146–59.

43. Alvarez M, and Casadevall A. Phagosome fusion and extrusion, and host cell survival following Cryptococcus neoformans phagocytosis by macrophages. Current Biology. 2006;16(2161–5.

44. Ma H, Croudace JE, Lammas DA, and May RC. Expulsion of live pathogenic yeast by macrophages. Curr Biol. 2006;16(21):2156–60.

45. Fu MS, and Casadevall A. Divalent metal cations potentiate the predatory capacity of amoeba for Cryptococcus neoformans. Appl Environ Microbiol. 2017.

46. Marietto-Gonçalves GA, and Grandi F. Are all psittacine birds carriers of Cryptococcus neoformans? Memorias do Instituto Oswaldo Cruz. 2011;106(6):781; author reply -2.

47. Nosanchuk JD, Mednick A, Shi L, and Casadevall A. Experimental murine cryptococcal infection results in contamination of bedding with Cryptococcus neoformans. Contemporary topics in laboratory animal science. 2003;42(4):9–12.

48. Lin J, Zhao Y, Ferraro AR, Yang E, Lewis ZA, and Lin X. Transcription factor Znf2 coordinates with the chromatin remodeling SWI/SNF complex to regulate cryptococcal cellular differentiation. Communications biology. 2019;2(412.

49. Jian Y, Liu Z, Wang H, Chen Y, Yin Y, Zhao Y, and Ma Z. Interplay of two transcription factors for recruitment of the chromatin remodeling complex modulates fungal nitrosative stress response. Nature communications. 2021;12(1):2576.

50. Yu CH, Chen Y, Desjardins CA, Tenor JL, Toffaletti DL, Giamberardino C, Litvintseva A, Perfect JR, and Cuomo CA. Landscape of gene expression variation of natural isolates of Cryptococcus neoformans in response to biologically relevant stresses. Microbial genomics. 2020;6(1).

51. Yu CH, Sephton-Clark P, Tenor JL, Toffaletti DL, Giamberardino C, Haverkamp M, Cuomo CA, and Perfect JR. Gene Expression of Diverse Cryptococcus Isolates during Infection of the Human Central Nervous System. mBio. 2021;12(6):e0231321.

52. Chen Y, Farrer RA, Giamberardino C, Sakthikumar S, Jones A, Yang T, Tenor JL, Wagih O, Van Wyk M, Govender NP, et al. Microevolution of Serial Clinical Isolates of Cryptococcus neoformans var. grubii and C. gattii. MBio. 2017;8(2).

53. Gusa A, Williams JD, Cho JE, Averette AF, Sun S, Shouse EM, Heitman J, Alspaugh JA, and Jinks-Robertson S. Transposon mobilization in the human fungal pathogen Cryptococcus is mutagenic during infection and promotes drug resistance in vitro. Proceedings of the National Academy of Sciences of the United States of America. 2020;117(18):9973–80.

54. Feldmesser M, Kress Y, Novikoff P, and Casadevall A. Cryptococcus neoformans is a facultative intracellular pathogen in murine pulmonary infection. Infect Immun. 2000;68(7):4225–37.

55. Wiseman H, and Halliwell B. Damage to DNA by reactive oxygen and nitrogen species: role in inflammatory disease and progression to cancer. The Biochemical journal. 1996;313 (Pt 1)(Pt 1):17–29.

56. Ene IV, Farrer RA, Hirakawa MP, Agwamba K, Cuomo CA, and Bennett RJ. Global analysis of mutations driving microevolution of a heterozygous diploid fungal pathogen. Proceedings of the National Academy of Sciences of the United States of America. 2018;115(37):E8688–e97.

57. Fu MS, Liporagi-Lopes LC, Dos Santos SRJ, Tenor JL, Perfect JR, Cuomo CA, and Casadevall A. Amoeba Predation of Cryptococcus neoformans Results in Pleiotropic Changes to Traits Associated with Virulence. mBio. 2021;12(2).

58. Martínez-Díaz R, Herrera S, Castro A, and Ponce F. Entamoeba sp.(sarcomastigophora: Endamoebidae) from ostriches (struthio camelus)(Aves: Struthionidae). Veterinary Parasitology. 2000;92(3):173–9.

59. Suthers HB. Ground-feeding migratory songbirds as cellular slime mold distribution vectors. Oecologia. 1985;65(4):526–30.

60. Litvintseva AP, and Mitchell TG. Most environmental isolates of Cryptococcus neoformans var. grubii (serotype A) are not lethal for mice. Infect Immun. 2009;77(8):3188–95.

61. Rivera J, Feldmesser M, Cammer M, and Casadevall A. Organ-dependent variation of capsule thickness in Cryptococcus neoformans during experimental murine infection. Infect Immun. 1998;66(5027–30.

62. Fernandes KE, Brockway A, Haverkamp M, Cuomo CA, van Ogtrop F, Perfect JR, and Carter DA. Phenotypic Variability Correlates with Clinical Outcome in Cryptococcus Isolates Obtained from Botswanan HIV/AIDS Patients. mBio. 2018;9(5).

63. Guerrero A, Jain N, Wang X, and Fries BC. Cryptococcus neoformans variants generated by phenotypic switching differ in virulence through effects on macrophage activation. Infect Immun. 2010;78(3):1049–57.

64. Polacheck I, Platt Y, and Aronovitch J. Catecholamines and virulence of Cryptococcus neoformans. Infect Immun. 1990;58(2919–22.

65. Goldman D, Lee SC, and Casadevall A. Pathogenesis of pulmonary Cryptococcus neoformans infection in the rat. Infect Immun. 1994;62(4755–61.

66. Goldman DL, Casadevall A, Cho Y, and Lee SC. Cryptococcus neoformans meningitis in the rat. Lab Invest. 1996;75(759–70.

67. Huffnagle GB, Yates JL, and Lipscomb MF. Immunity to pulmonary Cryptococcus neoformans infection requires both CD4+ and CD8+ T cells. J Exp Med. 1991;173(793–800.

68. Lagrou K, Van Eldere J, Keuleers S, Hagen F, Merckx R, Verhaegen J, Peetermans WE, and Boekhout T. Zoonotic transmission of Cryptococcus neoformans from a magpie to an immunocompetent patient. Journal of internal medicine. 2005;257(4):385–8.

69. Fries BC, Chen F, Currie BP, and Casadevall A. Karyotype instability in Cryptococcus neoformans infection. J Clin Microbiol. 1996;34(1531–4.

70. Forche A, Cromie G, Gerstein AC, Solis NV, Pisithkul T, Srifa W, Jeffery E, Abbey D, Filler SG, Dudley AM, et al. Rapid Phenotypic and Genotypic Diversification After Exposure to the Oral Host Niche in Candida albicans. Genetics. 2018;209(3):725–41.

71. Casadevall A, and Pirofski LA. Benefits and Costs of Animal Virulence for Microbes. mBio. 2019;10(3).

