## Supplementary Figures for "Human and murine *Cryptococcus neoformans* infection selects for common genomic changes in an environmental isolate"

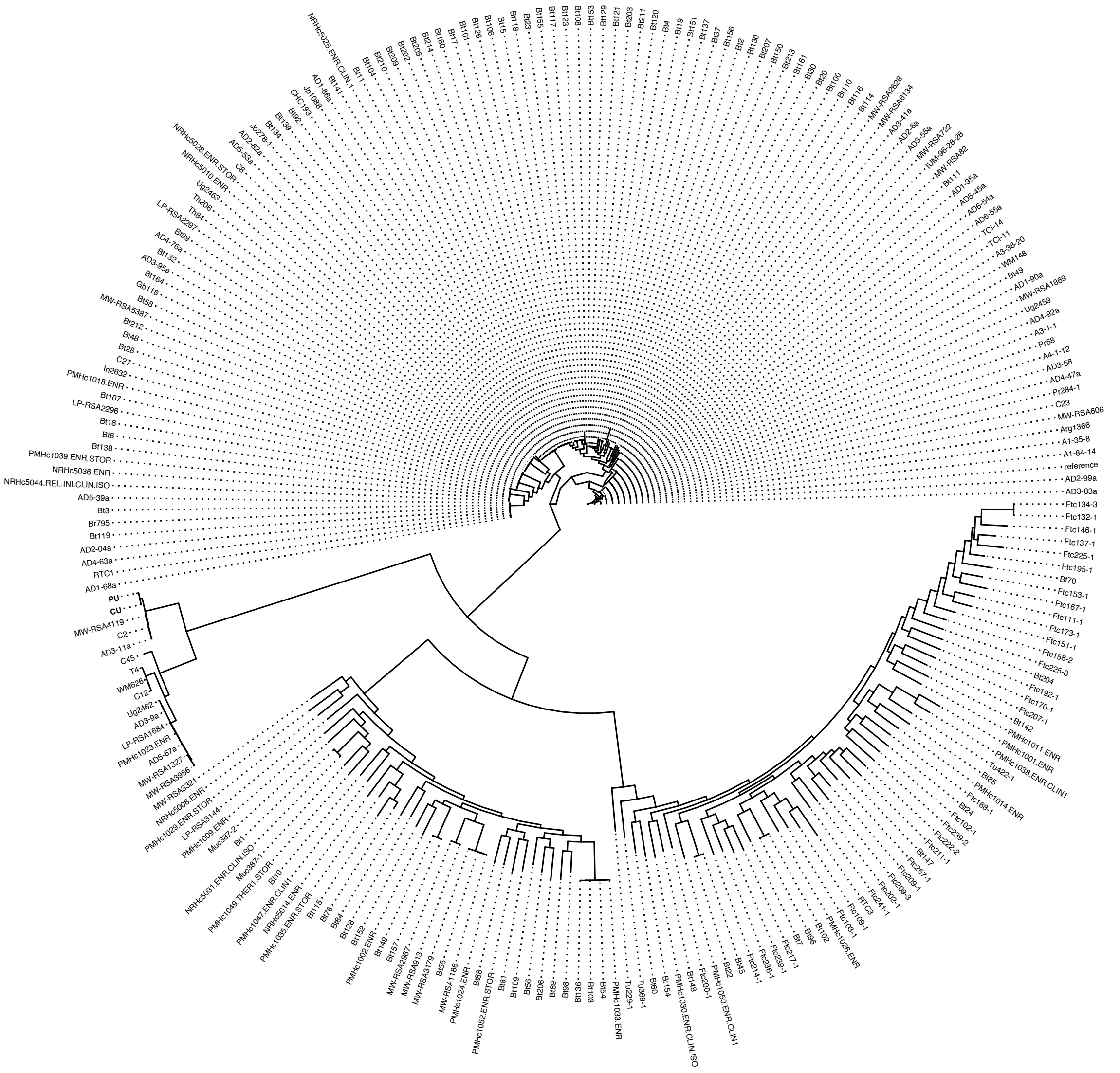

**Supplementary Figure 1. Cockatoo and patient strain phylogeny.** Maximum likelihood phylogeny of Desjardins et al. (34) isolates with the cockatoo (CU) and patient (PU) strains represented within the VNII lineage. All lineages show 100% bootstrap support.

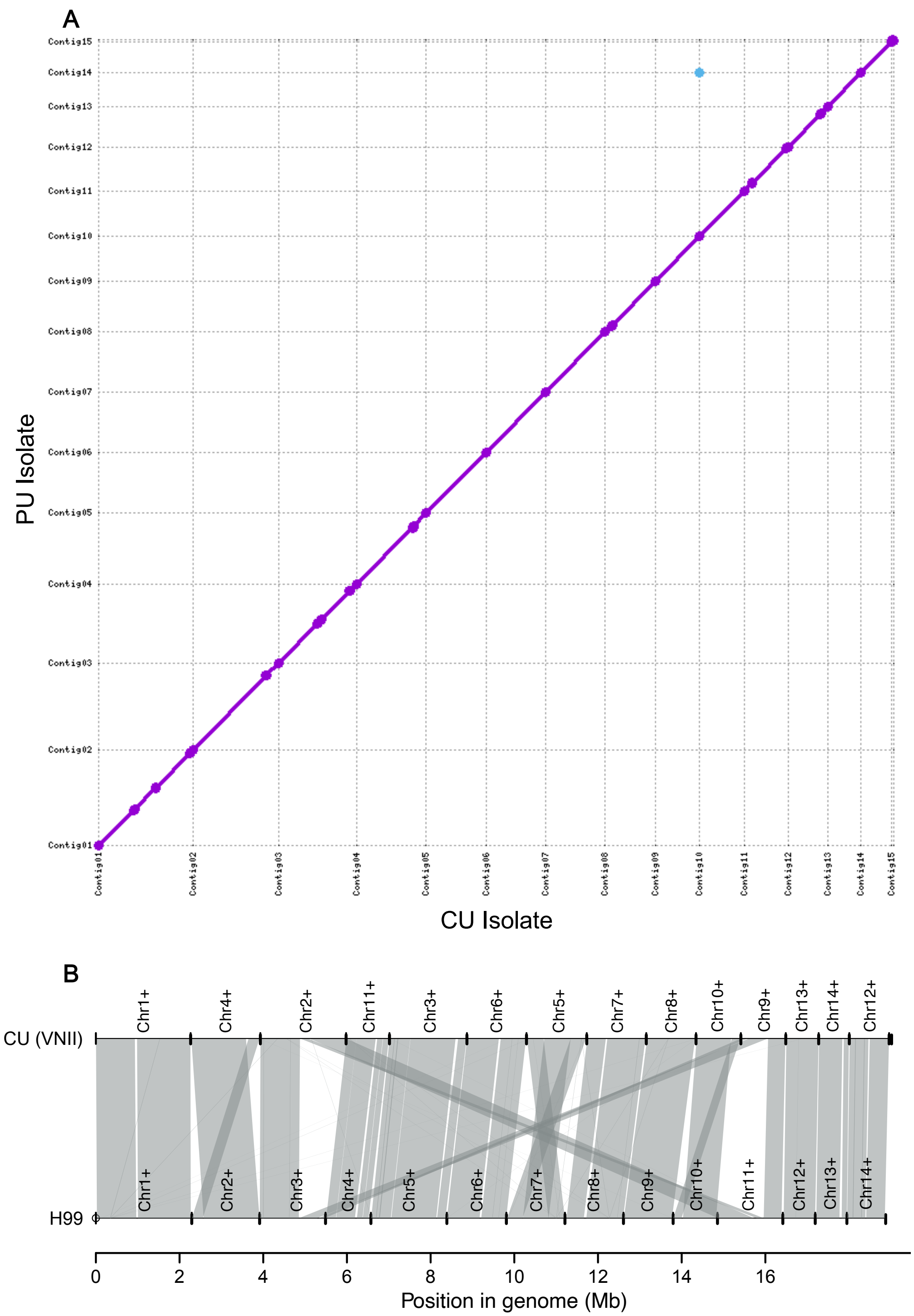


**Supplementary Figure 2. Synteny between genome assemblies.** (A) CU and PU genome assembly alignments. (B) Synteny between CU and *C. neoformans* H99 genome assemblies.

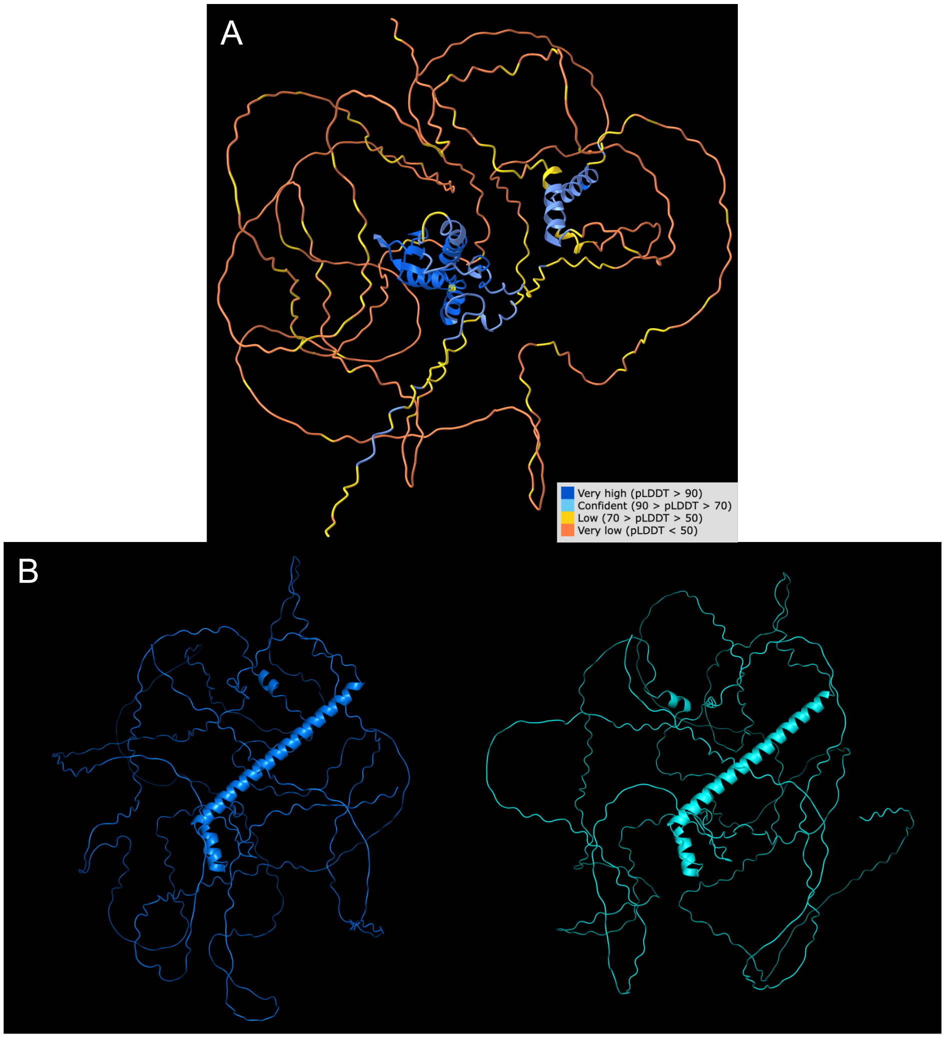


C

1 MAGSETSSPTDFAFEPPALPSGDAAFTCTPYKPEPLKPTFASPTGLSPPTGSFTPAANAA
61 PGHSYFPDSYNYGVSSASSFSSASASATGGSFSSVDNLPGISKNFVRPNLADTRRPATAG
121 GALQSRSPYSNFMMSSKNGMSGSMSQDAQKLGERGDMRRPEGTIEEGNEMLGLGGSTEED
181 GDEQGPQNVSPENENTVNPTLFPANRRSSAPQYNIPGSNWGQFPPSSSHDASNSSLGLPS
241 APAHMPMGYLQQQQAAHHNLNTQNRPPAFAGRPQTSDGLPSYTNYHGAVTLPSAQSIARQ
301 IPGMTDTTSFFHHSSGSAHLPFRDDRHFPSTNFPGDRAFTFDPLRSSLPPSFPGQIRASY
361 APPPPVPTAGSDSGGSSNNTNSTNDQMQFMSLSTGHGQKKRPRRRYEEIERLYPCGWNGC
421 EKSYGTLNHLNAHVMTQKHGEKRLPAEFKEMRKAWRKKKREHASAQASSQYMTNAAAWQQ
481 NYQRLSFASTSSAPDSDWDRRESSTSTVSVSTDGRPSFSYPTNYSWGVPPQGMPFQAGMM
541 SVDSRPSTSSSNVSSVSMDGTFISNQPTSASTSVMSMGPPTYLYLSGPPSLPGGVVSRRP
601 SAPQHVPMPPSNPVDGFRPADGDHPTPTMGNPFPTGPSGAGVVAGAGAGAGAAGQAGKAL
661 NFSTLTSPMKGSSGDFGSQFAFTR

**Supplementary Figure 3. Predicted structures for LQVO5_000317 and LQVO5_004463.** (A) Structure of LQVO5_000317, as predicted by AlphaFold2, with prediction confidence indicated by ribbon color. (B) Structure of CNAG_05940 (left) and the extended LQVO5_004463 (right) protein, as predicted by AlphaFold2. (C) CNAG_05940 protein sequence with intrinsically disordered region's (>40 residues) underlined, Ser/Thr regions highlighted and colored by predicted glycosylation state (red = glycosylated, green = un-glycosylated), and residues which differ in the extended LQVO5_004463 protein bolded (P>S and A>T).

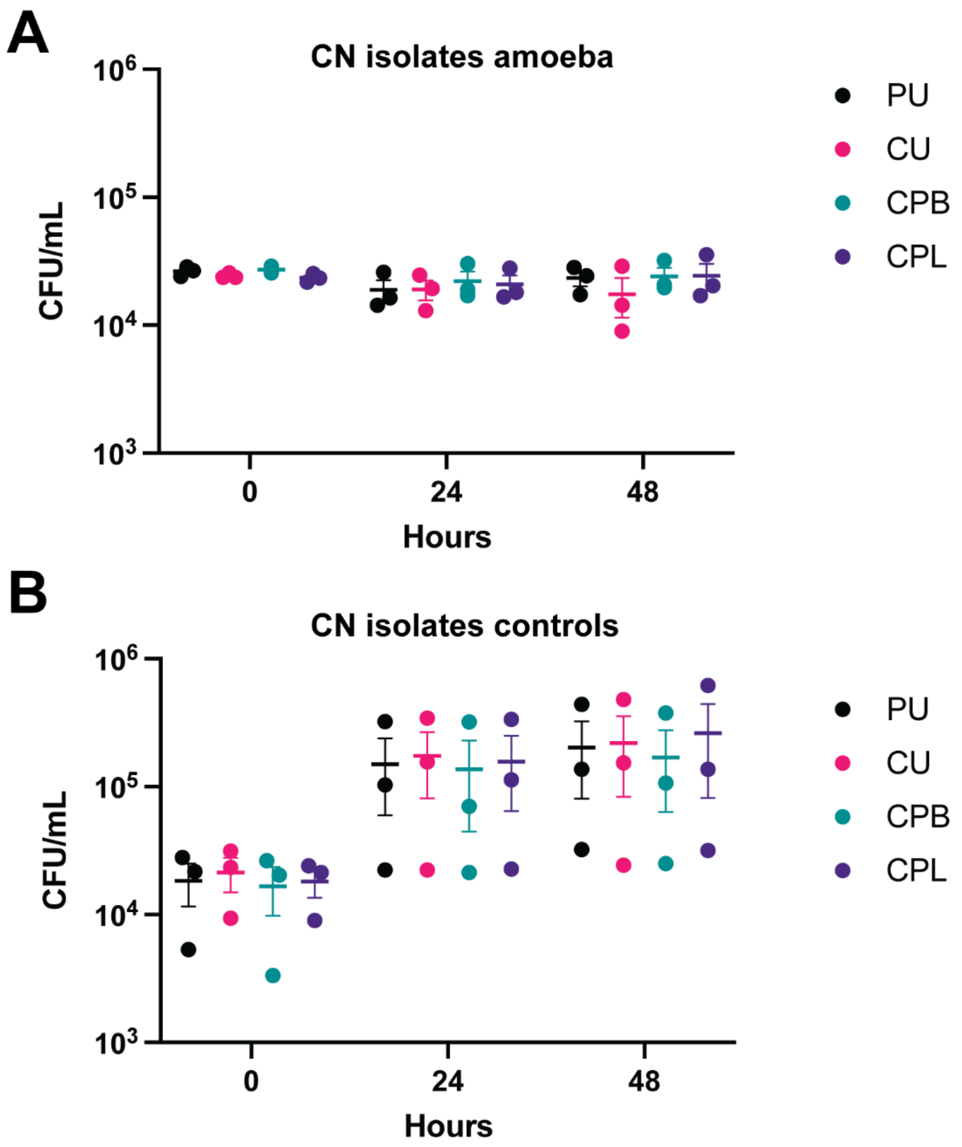


**Supplementary Figure 4. Amoeba interactions with *C. neoformans* strains and isolates.** Survival of *C. neoformans* strains and isolates after incubation with *A. castellanii*. Fungal survival was determined by CFU counts on YPD agar after 2 days of growth at 30 °C. (A) CFUs of *C. neoformans* strains and isolates incubated with *A. castellanii* for 0, 24, or 48 hours at 25 °C. (B) CFUs of *C. neoformans* strains and isolates incubated in 1x PBS for 0, 24, or 48 hours at 25 °C. Each data point represents an independent experiment.
